# An extra-genital cell population contributes to urethra closure during mouse penis development

**DOI:** 10.1101/2023.11.09.564741

**Authors:** Ciro Maurizio Amato, Xin Xu, Humphrey Hung-Chang Yao

## Abstract

Hypospadias, or incomplete closure of the urethra along the penis, is the second most common birth defect in the United States. We discovered a population of extra- genital mesenchymal cells that are essential for proper penile urethra closure in mouse embryos. This extra-genital population first appeared in the mesenchyme posterior to the hindlimb of the fetus after the onset of penis formation. These extra-genital cells, which transiently express a lineage marker *Nr5a1*, migrated centrally and colonized the penis bilateral to the urethra epithelium. Removal of the *Nr5a1+* extra-genital cells, using a cell-type specific ablation model, resulted in severe hypospadias. The absence of extra-genital cells had the most significant impacts on another mesenchymal cells, the peri-urethra that were immediately adjacent to the *Nr5a1+* extra-genital cells. Single cell mRNA sequencing revealed that the extra-genital cells extensively interact with the peri-urethra, particularly through Neuregulin 1, an epidermal Growth Factor (EGF) ligand. Disruption of Neuregulin 1 signaling in the *ex-vivo* slice culture system led to failure of urethra closure, recapitulating the phenotypes of extra-genital cell ablation. These results demonstrate that the *Nr5a1+* extra-genital mesenchymal cells from outside of the fetal penis are indispensable for urethra closure through their interaction with the peri-urethra mesenchymal cells. This discovery provides a new entry point to understand the biology of penis formation and potential causes of hypospadias in humans.

## Introduction

Defects of the external genitalia are among the most common birth defects in the world (1). One such defect is hypospadias, where the urethra fails to close at the tip of the penis and instead, exits somewhere along the shaft. Approximately 1:125 male newborns are affected by hypospadias. Although hypospadias is common, its etiologies are largely unknown. Disruptions in androgen signaling are commonly associated with hypospadias in both humans and mice (2–4). Gene mutations in androgen receptor (*Ar*), 5-α reductase (*Srd5a2*), and steroidogenic factor 1 (*Nr5a1*) are linked to hypospadias formation (5). Although these predisposing mutations provide a few possible causes for hypospadias, most human hypospadias cases remain unexplained (6). Identifying the basic mechanisms underlying development of the fetal penis from non-human species such as mouse could provide critical information to understand how hypospadias occurs.

In the male embryo, urethra closure begins at embryonic day (E)14.5 in mice or post conceptional week (PCW) 9.5 in humans (7, 8). In this initial phase of urethra closure, the urethral epithelium is found medially, along the underside of the penis.

From the tip to the base of the penis, the urethral epithelium is externally visible as a contiguous epithelial sheet that connects to the skin of the presumptive penis. The urethra epithelium invaginates at the medial aspect and forms a canal at the base of the penis. The immediate neighbor to the invaginated urethra epithelium is a population of mesenchymal cells known as the peri-urethra cells, which are extensively involved in urethra closure (9). As the urethral epithelium begins to disconnect from the skin and form a tube, the bilateral peri-urethra cells move medially and fuse, or “close”, at the midline of the penis, eventually leading to the formation of the urethra tube (9). The process of urethral closure is facilitated by crosstalk between the peri-urethra population and the urethral epithelium, and involves ligands such as sonic hedgehog (SHH), WNT, fibroblast growth factors (FGFs) and bone morphogenic proteins (BMPs) (7, 10, 11).

Neighboring the peri-urethra cells is an additional group of mesenchymal cells, which compose the prepuce of the penis. The role of preputial mesenchymal cells in urethra closure has only been hypothesized (12). Based on anatomical studies on hypospadiac penises in humans and mice, the prepuce rarely fuses when the urethra is open. As the urethra closes, the prepuce follows the medial movement of the periurethral mesenchyme, eventually fusing at the midline of the penis. Signaling factors between the prepuce and surrounding penis cells are not known.

The fact that 70% of the human cases of hypospadias do not have a known cause highlights the need of developing models to understand the basic biology of penile development. In this study, we applied mouse genetic models, single cell sequencing, and ex-vivo culture systems to discover cell types of the penis and the interactions among these cells in facilitating proper urethra closure.

## Results

### Identification of a transcriptionally distinct cell population in the fetal penis that is derived outside the external genitalia

Male external genitalia require androgens, particularly dihydrotestosterone, for its sex-specific morphogenesis. The external genitalia express 5α-reductase, the enzyme that converts testis-derived testosterone into dihydrotestosterone, a potent form of androgen that masculinizes the external genitalia (13, 14). Loss-of-function mutations of the gene encoding 5α-reductase (*SRD5A2*) resulted in feminization of the external genitalia and severe hypospadias in XY humans (15). Despite its critical functions in penis development, how 5α-reductase expression is activated in the fetal external genitalia has yet to be defined. The expression of many steroidogenic enzymes in gonads and adrenals are induced by the orphan nuclear receptor Steroidogenic factor 1 or NR5A1, the master regulator of steroidogenesis (16, 17). We therefore hypothesized that NR5A1 could be responsible for 5α-reductase expression in fetal external genitalia. To visualize cells that express *Nr5a1*, we developed the *Nr5a1-Cre^tg/-^*;*Rosa-LSL-tdTomato^+/f^* mouse model where *Nr5a1*+ cells are permanently marked with the fluorescent protein tdTomato (or *Nr5a1^tdTomato^+* cells henceforth). We found that at embryonic day 11.5 or E11.5, one day after the onset of penis formation in XY mouse embryos, no *Nr5a1^tdTomato^*^+^ cells were observed in any part of the posterior embryo including the external genitalia (**Supplemental Figure 1)**. However, one day later at E12.5, bilateral clusters of *Nr5a1^tdTomato^*^+^ cells appeared posterior to the hindlimbs in the inguinal region outside of the penis (**Figure 1A**). At E14.5 the *Nr5a1^tdTomato+^* cell clusters extended from the inguinal region to bilateral posterior sides of the penis (**Figure 1B**).

**Figure 1.**
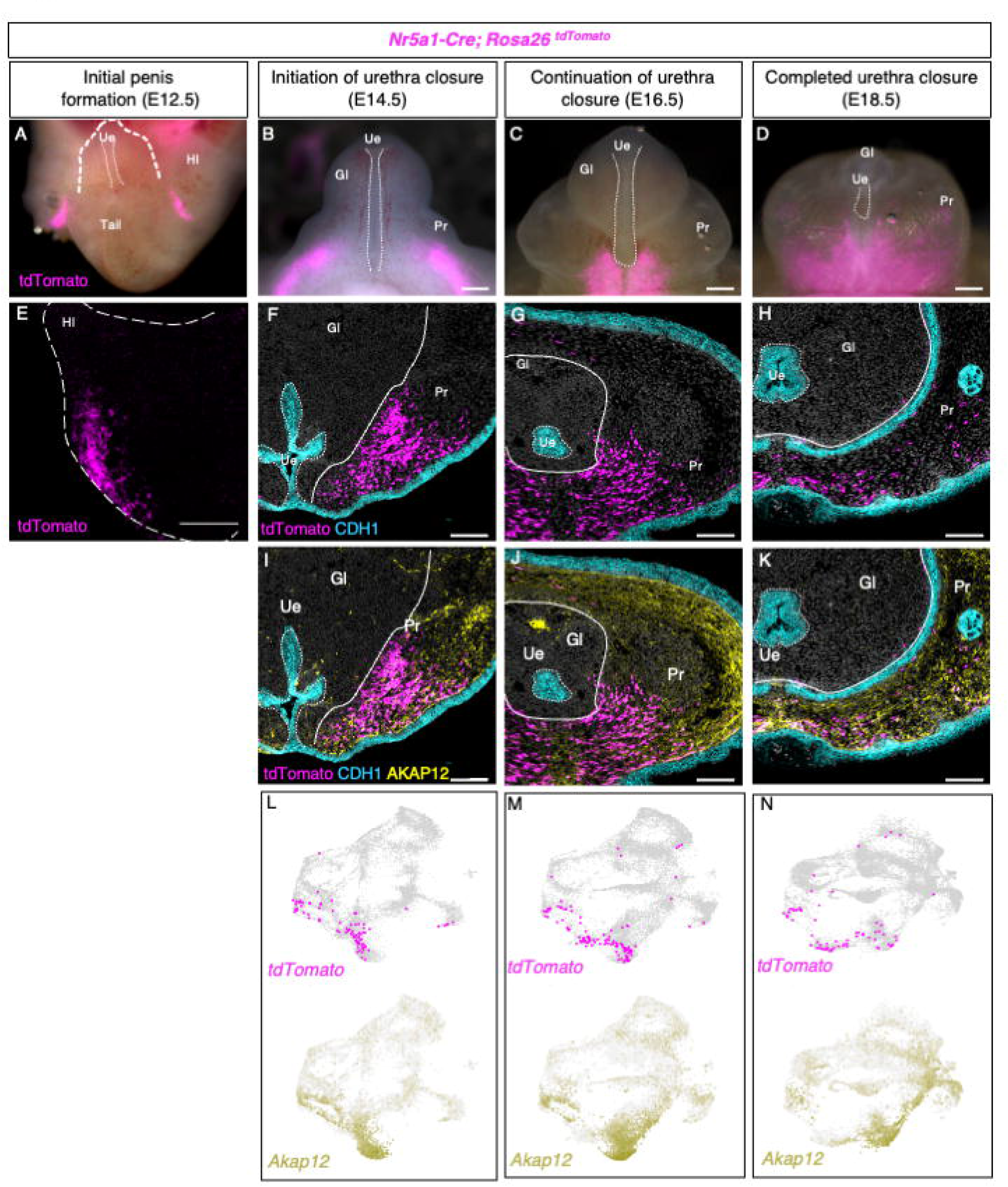
Contribution of hindlimb-derived *Nr5a1^tdTomato+^* cells to the fetal penis. **(A-D)** Whole mount images of external genitalia from E12.5-E18.5 XY mouse embryos. *Nr5a1-Cre; Rosa26^tdTomato^* mice were used to label hindlimb cells in magenta. White dotted lines in A-D mark the open urethra on the ventral aspect of penis. The bracketed line in A demarcates the fetal penis. **(E**) Lightsheet image of the right hindlimb of an E12.5 mouse embryo, *Nr5a1^tdTomato+^* cells are labeled in magenta. **(F-H**) Immunofluorescent staining of penis cross-sections at E14.5, E16.5, and E18.5 with magenta indicating cells with endogenous tdTomato fluorescence and blue labeling the epithelium (CDH1). (**I-K**) Overlay of prepuce marker AKAP12 from the same sections in E-F. (**L-N**) Single cell mRNA UMAP displaying the expression of tdTomato (red) and AKAP12 (yellow). **A-E** scale bars = 200µm and **F-K** scale bar = 100µm. Hl=Hindlimb, Gl = Glans, Pr= Prepuce, and Ue = urethra.

As the urethra continues to close (E16.5-18.5), the *Nr5a1^tdTomato+^* cells moved medially and eventually became confluent along the proximal-ventral aspect of the penis (**Figure 1C & D**).

To investigate the identity and location of *Nr5a1^tdTomato+^*cells, we performed immunofluorescence for tdTomato, CDH1 (a marker for the epithelium of the urethra and epidermis), and AKAP12 (a marker for the general prepuce). We also analyzed the single cell mRNA sequencing data that we published (18). In the wholemount E12.5 XY embryos, *Nr5a1*^tdTomato+^ cells were not present in the external genitalia and instead, were found under the epidermis of the inguinal region of the embryo (**Figure 1E** and **Supplemental Movie 1)**. At E14.5 onwards, the *Nr5a1*^tdTomato+^ cells became a part of the preputial swellings based on the partial colocalization of the *Nr5a1^tdTomato^* fluorescence with the general prepuce marker, AKAP12, in both the histological sections and single cell mRNA sequencing data (**Figure 1I-N**). At E14.5 and E16.5, ∼76% and 65% of *Nr5a1^tdTomato^*cells express *Akap12.* By E18.5, 30% of *Nr5a1^tdTomato^* cells express *Akap12*, suggesting that *Nr5a1^tdTomato^* cells are heterogenous and diverge from the general prepuce cell population. Throughout the rest of embryonic development, the *Nr5a1*^tdTomato+^ cells remained in the prepuce and eventually became a single, confluent cell population along the ventral aspect of the penis, where the urethra once was located (**Figure 1F-K**). *Nr5a1^tdTomato+^* cells expressed distinct sets of genes, including *Myocd, Grem1, Scube2 and* others (**Supplemental Figure 2** and **Supplemental Data 1**). These genes are commonly associated with myofibroblast and smooth muscle cell types.

The *Nr5a1-Cre^tg/+^*;*Rosa-LSL-tdTomato^+/f^*model permanently labels the cells as soon as they express *Nr5a1*. To profile the active expression of *Nr5a1* and determine whether *Nr5a1* is expressed in the genitalia, we examined the external genitalia using an additional *Nr5a1* reporter, *Nr5a1-GFP* (19). In this model, the GFP reporter was under the control of *Nr5a1* regulatory sequences and represented the active expression of *Nr5a1* at the time of examination. An identical pattern of *Nr5a1^tdTomato+^* cell population was observed at E12.5 outside of the penis (**Supplemental Figure 3A)**. However, in contrast to the *Nr5a1-Cre^tg/+^*;*Rosa-LSL-tdTomato^+/f^* lineage tracing model, *Nr5a1-GFP* expression was not found within the penis at any stage after E13.5 (**Supplemental Figure 3B**). This observation was also corroborated by our published single cell mRNA sequencing dataset, which showed that *Nr5a1* gene expression was detected in very few cells (2–5) throughout penis development (**Supplemental Figure 3C**).

We next investigated whether *Nr5a1* is functionally important for penis development by controlling the expression of *Srd5a2*. We developed a cell specific knockout of *Nr5a1* with the *Isl1^cre/+^* mice that target hindlimb and external genitalia (20). We first examined the *Isl1Cre* activity in our hand by crossing the *Isl1^cre/+^* mice to the *Rosa-tdTomato* reporter mice. We found that the *Isl1Cre* activity occurred at E10.5 prior to *Nr5a1* expression in the hindlimbs of the embryo (**Supplemental Figure 4A and B**). By postnatal day 0, almost the entire penis was positive for *Isl1Cre* activity (**Supplemental Figure 4C and D**), indicating its feasibility to target *Nr5a1* deletion.

When the *Nr5a1* gene was inactivated in the hindlimbs and genitalia (*Isl1^cre/+^*; *Nr5a1^f/-^),* the penis developed normally with the prepuce fusing along the ventral aspect of the penis and the urethra exiting at the tip (**Supplemental Figure 5A and B**). Additionally, inactivation of *Nr5a1* in the penis did not affect *Srd5a2* expression (**Supplemental figure 5C and D**). Together these data reveal that *Nr5a1* is a lineage identifier that transiently marks a distinct progenitor population derived from outside of the embryonic penis. These extra-genital derived *Nr5a1^tdTomato+^* cells migrate centrally and eventually becomes a portion of the prepuce of the penis that is associated with urethra closure. The expression of *Nr5a1* did not have an impact on urethra closure or *Srd5a2* expression.

### The *Nr5a1*^tdTomato+^ cells actively migrate during urethra closure

To visualize the movement of the *Nr5a1*^tdTomato+^ cells during urethra closure, we developed an ex vivo penis slice culture system that enables us to observe the movement of *Nr5a1*^tdTomato+^ cells in real time. We collected external genitalia at E15.5, when urethra closure starts, and sliced them into 150 µm thick sections for live imaging. At the beginning of the culture (0 hour), the urethra was physically connected to the skin and was not a closed tube in the center of the penis (**Figure 2A**). The *Nr5a1*^tdTomato+^ cells at this time showed few signs of active cell migration (a lack of visible lamellipodia) and were bilateral to the central urethra (**Figure 2A and D; Supplemental movie 1**).

**Figure 2.**
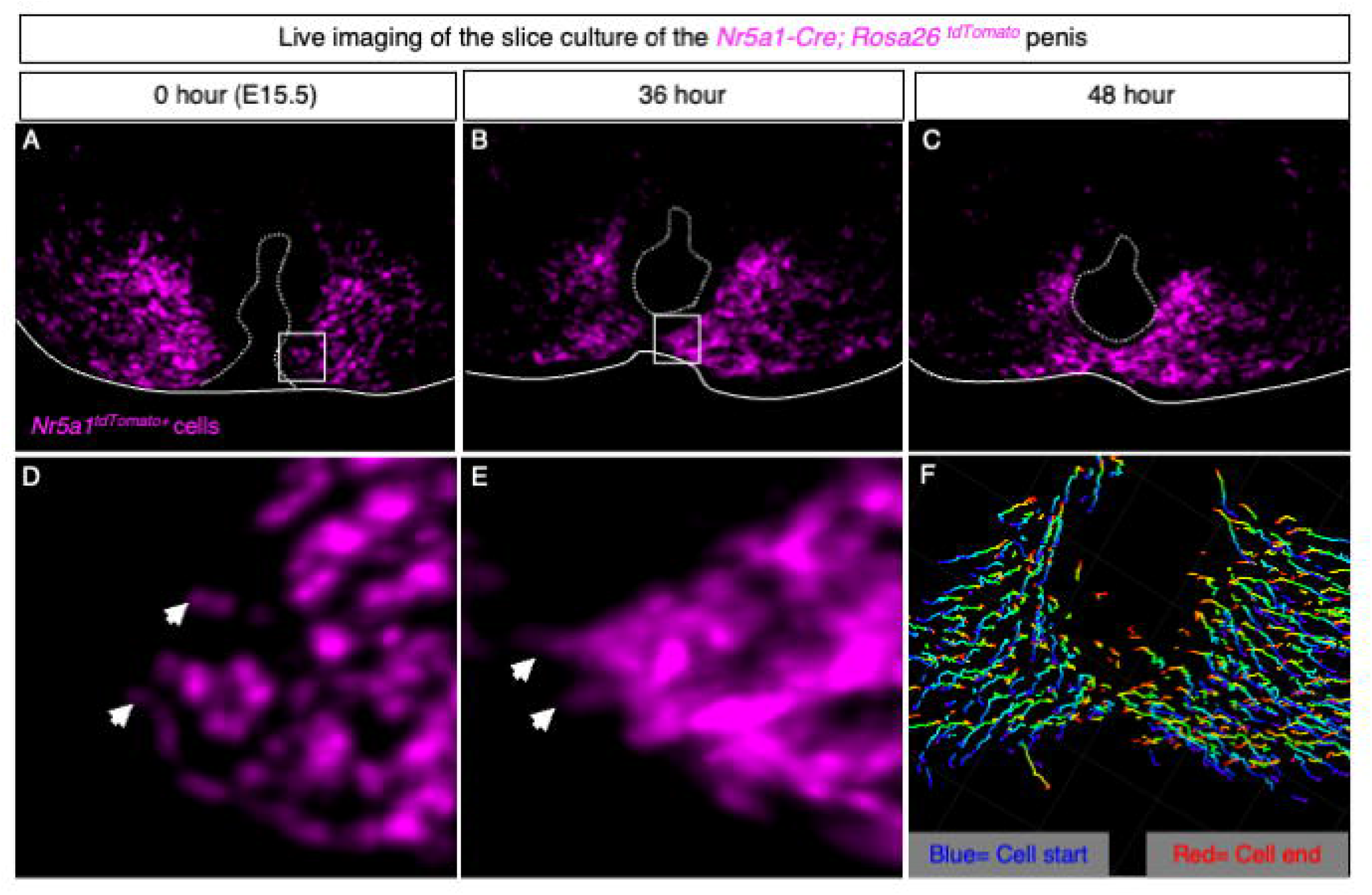
*Nr5a1^tdTomato+^* cells undergo extensive and active migration during urethra closure. (**A-C**) Fluorescent images of penis slice cultures from *Nr5a1-Cre; Rosa26^tdTomato^* embryos. Dotted lines represent the urethra and solid lines outline the penis slice. White boxes in **A** and **B** represent magnified regions in **D** and **E**. White arrowheads designate filopodia in the same cell at 0 and 36 hrs. (**F**) Dragon tail tracks generated by Imaris to mark the cell migratory paths. Blue colors represent the cell position at 0 hr and red color is the position of the cells at 48 hr.

Thirty-six hours after culture, lamellopodia were observed at the leading edge of the *Nr5a1*^tdTomato+^ cell population, and the distance between the bilateral populations of *Nr5a1*^tdTomato+^ cells shortened (**Figure 2B and E, Supplemental movie 1**). By 48 hours, the *Nr5a1*^tdTomato+^ cells filled in the space ventrally where the urethra was once open, resulting in a closed urethra (**Figure 2C, Supplemental movie 1**). Upon investigating the migration tracks using the Imaris software, we found that the majority of the *Nr5a1*^tdTomato+^ cells migrate medially toward the urethra (**Figure 2F**).

### Loss of *Nr5a1*^tdTomato+^ cells results in severe hypospadias

To test if the *Nr5a1*^tdTomato+^ cells are essential for urethra closure, we ablated these cells using the diphtheria toxin cell ablation or *Rosa-LSL-DTA model* (21). In the presence of *Nr5a1-Cre^Tg^*^/+^, diphtheria toxin or DTA was induced, which halted translation and eventually caused cell death of the *Nr5a1*^tdTomato+^ cells. Ablation of *Nr5a1*^tdTomato+^ cells disrupted urethral closure as early as E16.5 (**Figure 3A** and **E**). At E17.5, when urethra closure had further progressed in the control embryos, the urethra of the *Nr5a1^tdTomato+^*cell ablated mice remained open (**Figure 3B** and **F).** By E18.5 and P0, hypospadias was clearly apparent in the *Nr5a1*^tdTomato+^ cell ablated mice with a failure of urethra closure (**Figure 3C, D, G,** and **H)**. Invariably, all *Nr5a1*^tdTomato+^ cell ablated male mice developed extremely severe cases of hypospadias and preputial hypoplasia (**Figure 3G** and **H**). In some cases, no preputial fusion occurred along the ventral aspect of the penis (22). Histological sections demonstrated that the urethra was open at the base of the *Nr5a1*^tdTomato+^ cell ablated penis in contrast to the control male where the urethra was completely closed (**Figure 3I and J**). We further confirmed that *Nr5a1^tdtomato+^*cell ablation sufficiently removed the majority of the *Nr5a1^tdTomato+^*cells in these hypospadiac penises (**Figure 3K** and **L**). Overall, our results demonstrate that ablation of *Nr5a1^tdtomato+^* cells resulted in severe hypospadias.

**Figure 3.**
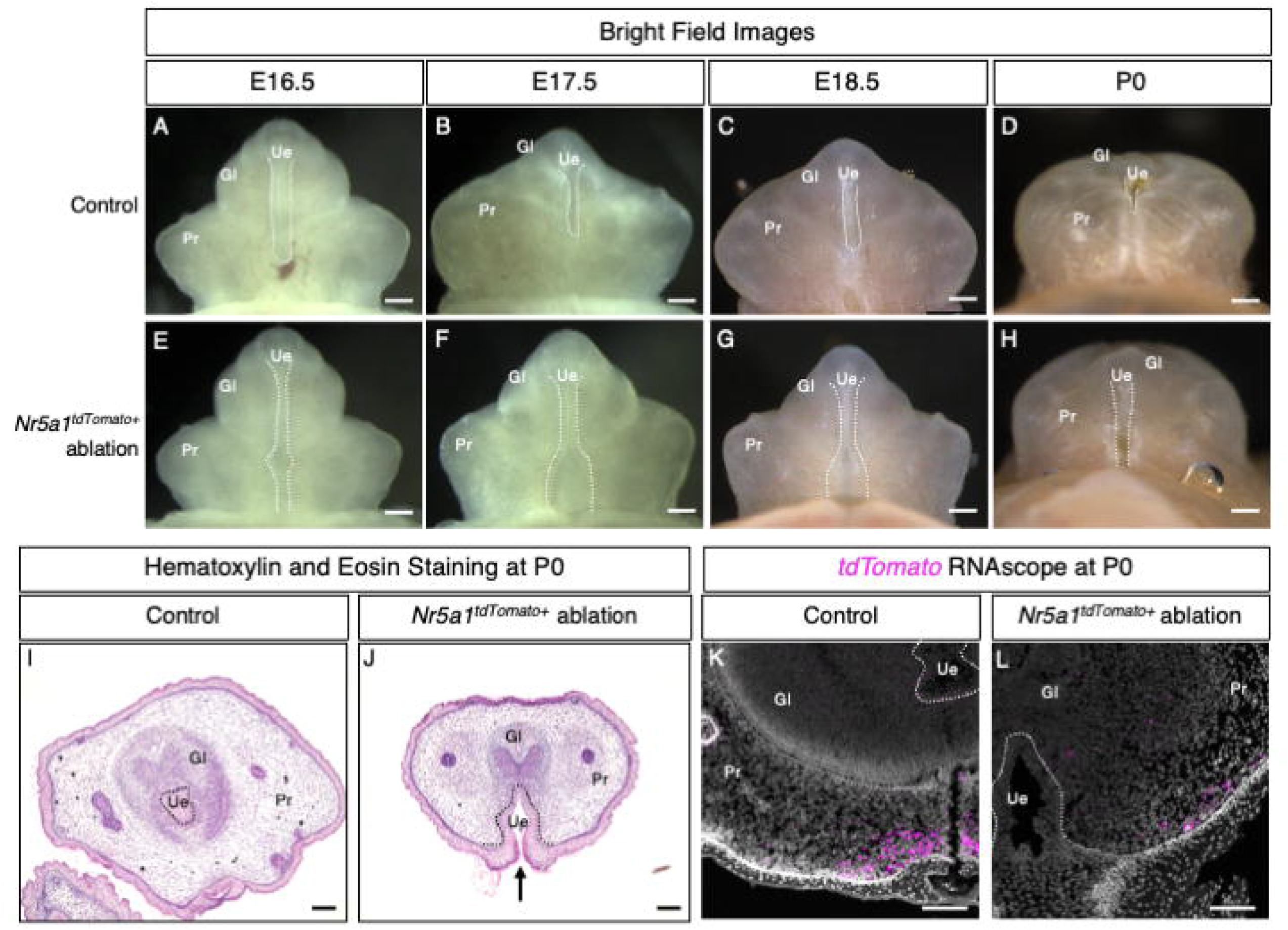
*Nr5a1^tdTomato+^* cells are required for urethra closure. (**A**-**H**) Whole mount images of control (**A-D**) and *Nr5a1^tdTomato+^* cell ablated (**E-H**) penises at E16.5, E17.5, E18.5 and P0. Dotted lines display regions where the urethra is open. **(I and J**) Hematoxylin and Eosin staining of penis sections from control and *Nr5a1^tdTomato+^* cell ablated embryos with the black dotted lines indicating the urethra and the black arrow indicating the open urethra. **(K** and **L**) RNAscope for *tdTomato* in control and *Nr5a1^tdTomato+^* cell ablated penis sections. Gl = Glans, Pr= Prepuce, and Ue = Urethra.

The *Nr5a1-Cre* induced ablation occurred in not only the penis, but also the gonads and adrenals. The size of the gonads in *Nr5a1^tdTomato+^* cell ablated embryos was drastically reduced and the adrenal glands could not be found in any of the embryos, demonstrating the efficiency of the cell ablation model (**Supplemental Figure 6A-C)**. However, without functional testes, androgen production would also be compromised in the embryo, leading to secondary impacts on the androgen-responsive organs such as the external genitalia. To exclude the possibility that the hypospadias phenotypes in the *Nr5a1^tdTomato+^* cell ablated penis was secondary to the degenerated testes, we supplemented the *Nr5a1^tdTomato+;^Rosa-DTA* embryos with 0.2 mg/10g of testosterone propionate during the critical window of urethra closure (E13.5-E18.5) by injecting the testosterone to the pregnant dam that carried these embryos. The dose of testosterone was based on previously published papers (2), which was sufficient to masculinize female embryos, using androgen-controlled anogenital distance as the indicator (**Supplemental Figure 6D and E)**. The *Nr5a1^tdTomato+^*cell ablated male embryos that were exposed to testosterone propionate still developed severe cases of hypospadias in 75% of the animals **(Supplemental Figure 6D)**. The partial rescue of the hypospadias phenotype is likely due to the variation of diptheria toxin efficiency between the different animals. These data indicate that some of the hypospadias phenotype due to the loss of testis-derived androgens, but mice do still develop severe hypospadias with the loss of *Nr5a1^tdTomato+^* cells.

Next, we investigated how the loss of *Nr5a1*^tdTomato^+ cells influence cell population dynamics in the penis. We conducted single cell mRNA sequencing on control (*Nr5a1-Cre^+/+^; RosaDTA^+/f^)* and *Nr5a1*^tdTomato+^ cell ablated (*Nr5a1-Cre^Tg/+^; RosaDTA^+/f^)* penises at the time of urethra closure (E16.5). The data were aligned, normalized, and unbiasedly clustered based on the transcriptome of each cell. We found extensive overlap between these datasets and our published single-cell datasets, confirming the reproducibility of our single cell analyses (18, 23). The unbiased cell clusters, or the composition of the cell populations, were similar between the control and *Nr5a1*^tdTomato+^ cell ablated penis; however, transcriptomic shifts in several cell populations were identified (**Figure 4A and B**). Markers of the *Nr5a1^tdTomato+^* cell population, *Grem1* and *Scube2* were significantly down-regulated, confirming the loss of *Nr5a1^tdTomato+^* cells (**Supplemental figure 7A and B**). Upon investigating the number of differentially expressed genes between the control and *Nr5a1^tdTomato+^* cell ablated samples for each penis cell population, we found that the corpus cavernosum, peri- urethra, ventral distal glans, preputial mesenchyme, dorsal distal glans and subdermal prepuce had the first (637), second (425), third (422), fourth (385), fifth (336), and sixth (217) most changed genes as a result of loss of *Nr5a1*^tdTomato+^ cells (**Figure 4C** and **Supplemental figure 8A**). The urethra epithelium had 33 genes differentially expressed and was one of the cell populations with the least amount of gene expression change due to *Nr5a1^tdTomato+^* cell ablation (**Figure 4C** and **Supplemental figure 8A**). These data indicates that removal of *Nr5a1*^tdTomato+^ cells have a global, and yet cell type-specific impact on the cell populations of the external genitalia.

**Figure 4.**
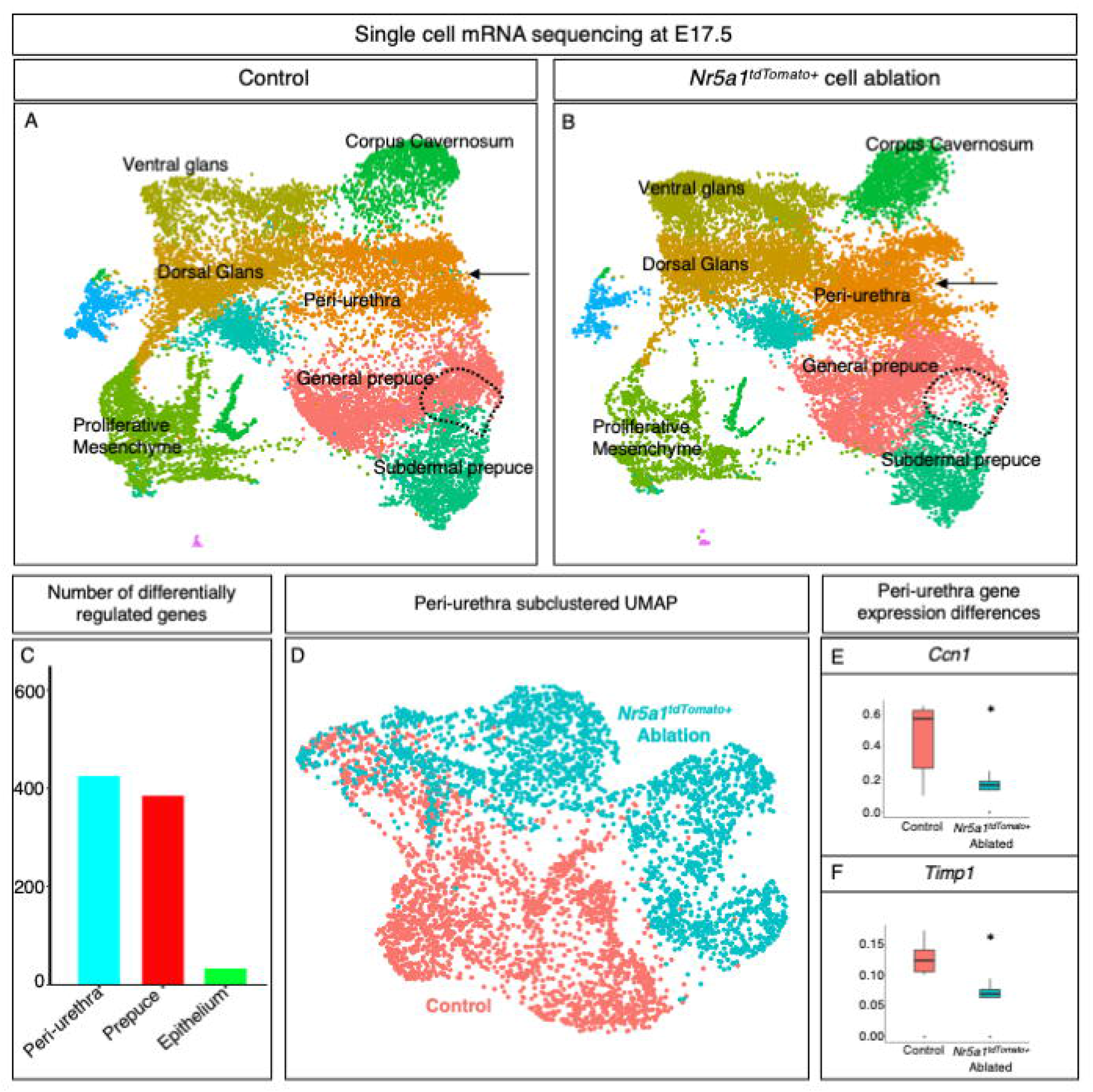
*Nr5a1^tdTomato+^* cell ablation induces transcriptomic changes in penis cell populations. **(A** and **B**) UMAP of control (**A**) and *Nr5a1^tdTomato+^* cell ablated (**B**) penises with colors representing unbiased clusters of cells. Black dotted line represents a region where *Nr5a1^tdTomato^*^+^ cells are lost in *Nr5a1^tdTomato+^* ablated penises and black arrows represent a shift in peri-urethra transcriptome. (**C)** Bar graph of differentially regulated genes in the peri-urethra, prepuce, and epithelium. (**D**) Reclustered UMAP of peri-urethral cells from control (red) and *Nr5a1^tdTomato+^* cell ablated penises (blue). (**E** and **F)** Significantly altered genes (*Ccn1* is a cell migration related gene and *Timp1* is an extracellular matrix modifier) in *Nr5a1^tdTomato+^* cell ablated peri-urethra. * represents p-value < 0.05.

The *Nr5a1^tdTomato+^* cells, which reside in the prepuce, are immediate neighbors to the peri-urethra mesenchymal cells, a cell population essential for urethra closure (9). To gain insight into how the peri-urethra cells were affected in the absence of *Nr5a1*^tdTomato+^ cells, we bioinformatically isolated these cells from the single cell dataset (**Figure 4A & B**), and re-clustered them (**Figure 4D**). The peri-urethra cells in the *Nr5a1*^tdTomato+^ cell ablated penis were significantly different from the control in their transcriptome with 654 differentially expressed genes (**Figure 4D**). Gene ontology analysis on the differentially expressed genes showed significant changes in pathways such as axonal guidance, hepatic fibrosis, and pulmonary fibrosis (**Supplemental figure 8B**). In the axonal guidance pathways, expression of several cell migration related genes (*Cnn1, Dner, Spock2, Nav3, Cxcl1,* and others) was altered, while the fibrotic pathways displayed strong changes in extracellular matrix gene expression (*Timp1, Itga4,* and *others)* (**Figure 4E** and **F and Supplemental figure 8C**). The loss of *Nr5a1*^tdTomato+^ cells have a significant impact on the peri-urethra cells, particularly the expression of extracellular matrix and cell migration genes in these cells.

### *Nr5a1^tdTomato+^* cells interact extensively with the neighboring peri-urethra mesenchyme

Urethra closure is facilitated by close communication between the urethra epithelium and its neighboring peri-urethra mesenchymal cells through signaling pathways such as WNT and SHH (24, 25). The *Nr5a1*^tdTomato+^ cells, which are directly bilateral to the peri-urethra mesenchyme, represent a newly identified cell population that could interact with urethra epithelium and peri-urethra mesenchyme to coordinate urethra closure (**Figure 5A-D**). To test this hypothesis, we identified potential interactions between secreted ligands and their receptors expressed by these three distinct cell types over the course of urethra closure by further analyzing our published single cell sequencing data with the CellPhoneDB v2.0 package (**Figure 5A**) (26). At the initiation of urethra closure (E14.5), extensive potential interactions (211 ligand-receptor pairs) were detected between the *Nr5a1*^tdTomato+^ cells (magenta) and the peri-urethra (yellow) mesenchyme (**Figure 5B & E).** Interactions between the epithelium (cyan) and either the peri-urethra or *Nr5a1*^tdTomato+^ cells were low to moderate (**Figure 5B & E**). Gene Ontology analyses of the ligands and receptor pairs between the *Nr5a1*^tdTomato+^ cells and peri-urethra revealed a significant enrichment of fibrotic (pulmonary fibrosis, hepatic fibrosis, and pulmonary healing) and cell migration (axonal guidance, ephrin signaling, and wound healing) related pathways. Similar relationships were found at both the continuation of urethra closure at E16.5 and completed urethra closure stages at E18.5 (**Figure 5F, G, I, J**). At these two stages, interaction between *Nr5a1*^tdTomato+^ and peri-urethra cells remained to be the strongest, although the number of ligand-receptor interactions decreased at E18.5 (n=127) (**Figure 5E-G**). Results from these analyses reveal that there is extensive communication between the *Nr5a1*^tdTomato+^ cells and the neighboring peri-urethra, and a moderate amount of signaling between *Nr5a1*^tdTomato+^ cells and urethral epithelial cells. The main interactions between the peri- urethra and *Nr5a1*^tdTomato+^ cells involve extracellular matrix modifications and cell migration pathways.

**Figure 5.**
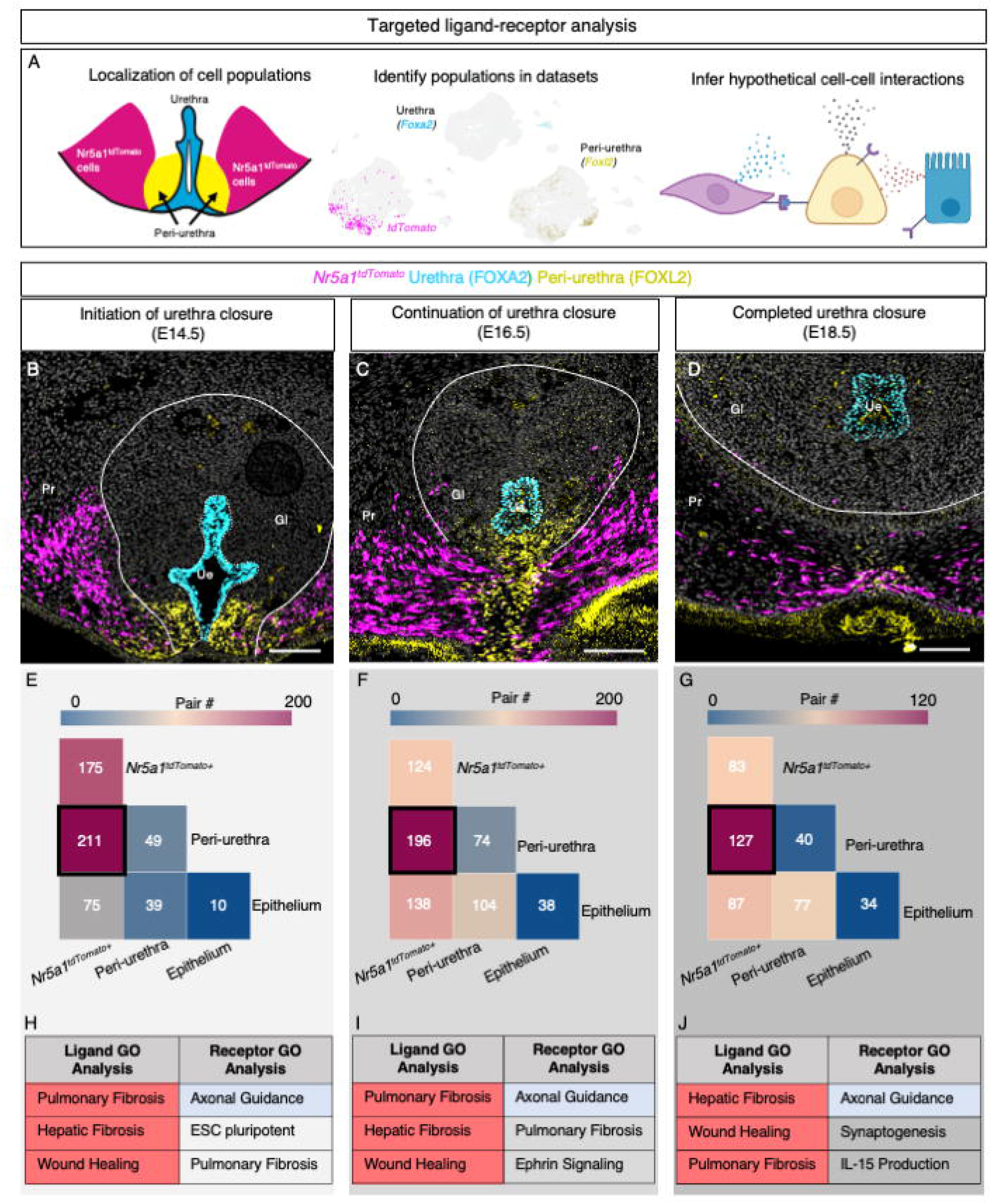
*Nr5a1^tdTomato+^* cells communicate extensively with neighboring mesenchymal cells. **(A)** Depiction of experimental design to identify potential ligand/receptor interactions. (**B-D)** Immunofluorescence of coronal penis sections at E14.5, E16.5, and E18.5, with FOXA2 (cyan for urethra), FOXL2 (yellow for peri- urethra), and tdTomato (magenta for *Nr5a1^tdTomato+^*cells). Scale bars in are 100µm. Gl = Glans, Pr= Prepuce, and Ue = Urethra. (**E-G**) Tile plots were used to present the numbers of ligand/receptor pairs with red = high number of ligand receptor pairs, and blue = low ligand receptor pairs. (**H-J**) GO analysis of ligands and receptors identified from the ligand receptor analysis within the peri-urethra cell population. Terms are listed in order of significance. Magenta shaded cells represent conserved processes for ligands, blue shaded cells represent conserved ontologies for receptors, and grey shaded cells represent ontologies not conserved between all stages.

### EGF paracrine signaling is essential for urethra closure and *Nr5a1*^tdTomato+^ cell migration

To identify the mechanism through which *Nr5a1*^tdTomato+^ cells facilitate urethra closure, we searched through the CellPhoneDB results to identify critical paracrine signaling factors from *Nr5a1*^tdTomato+^ cells (18). *Neuregulin-1 (Nrg1),* a member of the Epidermal growth factor or EGF family, was significantly enriched in the *Nr5a1*^tdTomato+^ cells compared to peri-urethra mesenchyme and urethra epithelium just before the initiation of urethra closure (E14.5) (**Figure 6A**). Other EGF members (*Egf, Tgfa, Areg, Btc,* and *Ereg)* had little to no expression in most of the cell populations (**Figure 6A**). Using RNAscope, we found considerable enrichment of *Nrg1* mRNA in the *Nr5a1*^tdTomato+^ cells in the penis (**Figure 6B and C**), supporting the single cell mRNAseq data.

**Figure 6.**
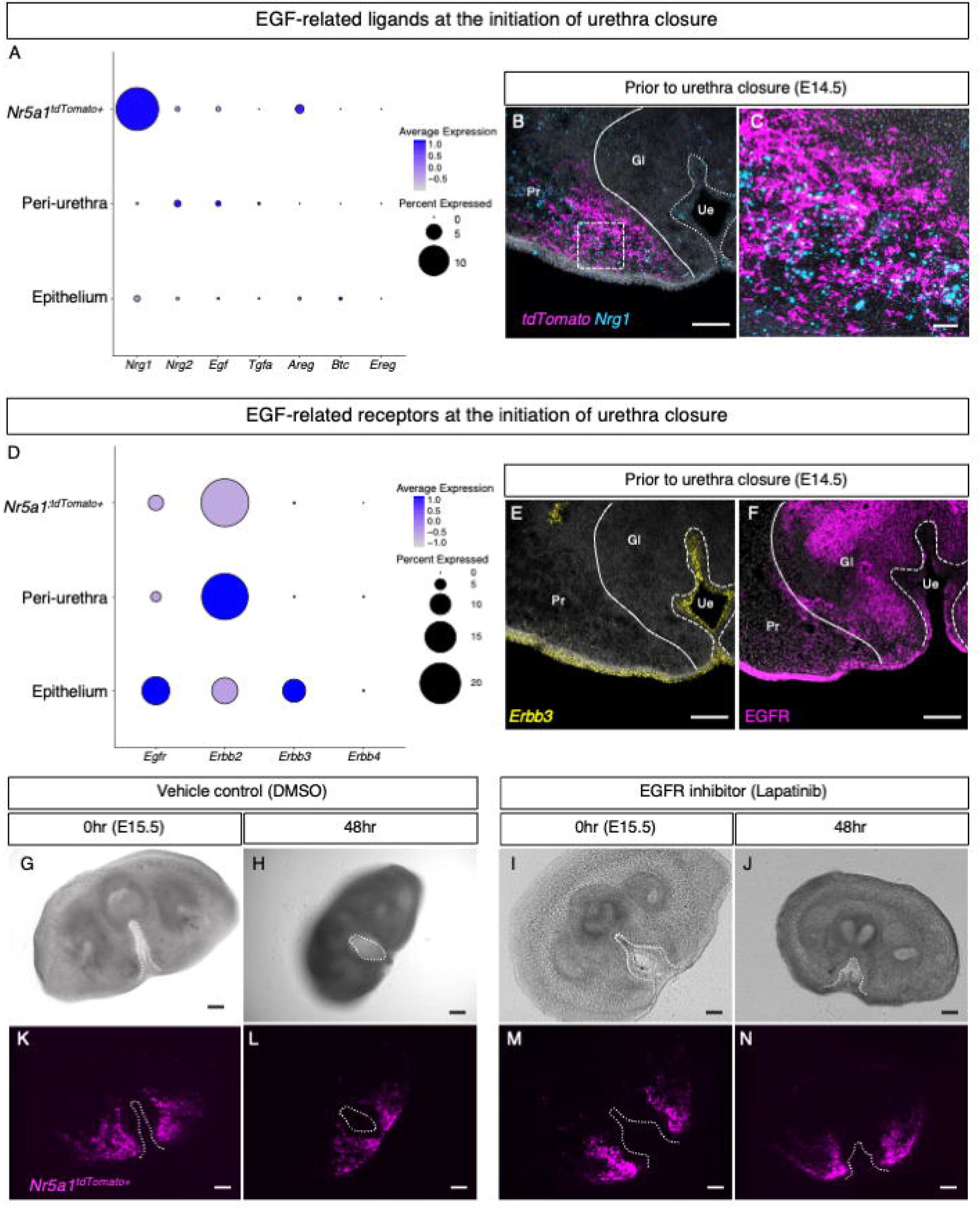
EGF signaling from *Nr5a1^tdtomato+^* cells as a critical pathway for urethra closure. (**A**) Dot plot of EGF-related ligand gene expression from single cell mRNA sequencing data of normal genitalia at E14.5. (**B and C)** low and high magnifications of RNAscope images of control penis sections at E14.5 stained for *Nrg1* (Cyan) and *tdTomato* (magenta). (**D**) Dot plot of EGF-related receptor gene expression from single cell mRNA sequencing data of normal genitalia at E14.5. (**E and F)** RNAscope and immunofluorescent images of control penis sections for *Erbb3 (*yellow) and EGFR (magenta) (**G-J**) Brightfield and (**K-N**) fluorescent images of control and EGFR inhibitor exposed slice culture sections at 0 and 48hrs. White dotted lines outline the urethra in all images. Scale bars in **B** and **E-N** are 100µm and **C** is 20µm. Gl = Glans, Pr= Prepuce, and Ue = Urethra.

NRG1 proteins signal through the receptor tyrosine kinases ERBB3 and ERBB4, and with less affinity to EGFR in several cell types (27). ERBB2 has no known ligand and mainly serves as a critical co-regulator for the other three EGF-related receptors (28). The peri-urethra mesenchyme had significant enrichment of *Egfr* and *Erbb2* receptors whereas the urethra epithelium expressed *Egfr, Erbb2,* and *Erbb3* (**Figure 6**). The *Nr5a1*^tdTomato+^ cells had some enrichment of *Egfr* and *Erbb2;* however, expression intensity of these genes was low. These data implicate both the peri-urethra and urethra epithelium as potential targets of *Nr5a1*^tdTomato+^ derived NRG1.

To determine whether EGF signaling through *Nr5a1^tdtomato+^*cell-derived NRG1 is essential for urethra closure, we took advantage of the slice culture system that we developed (**Figure 2**) and cultured the *Nr5a1^tdtomato+^*penis slices with Lapatinib, an EGFR and ERBB inhibitor, or di-methyl sulfoxide (DMSO) as the vehicle control (**Figure 6G-N**). Lapatinib specifically inhibits the phosphorylation and activation of the tyrosine kinases, EGFR and ERBB2 (29, 30). Penis slices exposed to DMSO underwent normal urethra closure with the *Nr5a1*^tdTomato+^ cells completing full migration (**Figure 6G, H, K, and L, Supplemental Movie 3**). On the other hand, Lapatinib inhibited *Nr5a1*^tdTomato+^ cell migration and led to failure of urethra closure (**Figure 6 I, J, M and N** and **Supplemental movie 4)**. These data show that the EGF signaling, likely activated by NRG1 from the *Nr5a1*+ mesenchymal cells, facilitates normal urethra closure.

## Discussion

### An extra-genital source of a unique cell type is essential for urethra closure

Formation of the penis requires the coordination of cells from unique developmental origins. Cells derived from the umbilicus and the tailgut undergo extensive outgrowth and differentiation to form the dorsal and ventral aspects of the genital tubercle, respectively. Here, we identify that a transcriptionally distinct group of cells, originating from the inguinal region of the embryo, contributes to normal penis development, particularly the closure of urethra (**Figure 7**). Once these extra-genital cells enter the penis, they undergo extensive migration and secrete paracrine factors that support normal urethra closure (**Figure 7**).

**Figure 7.**
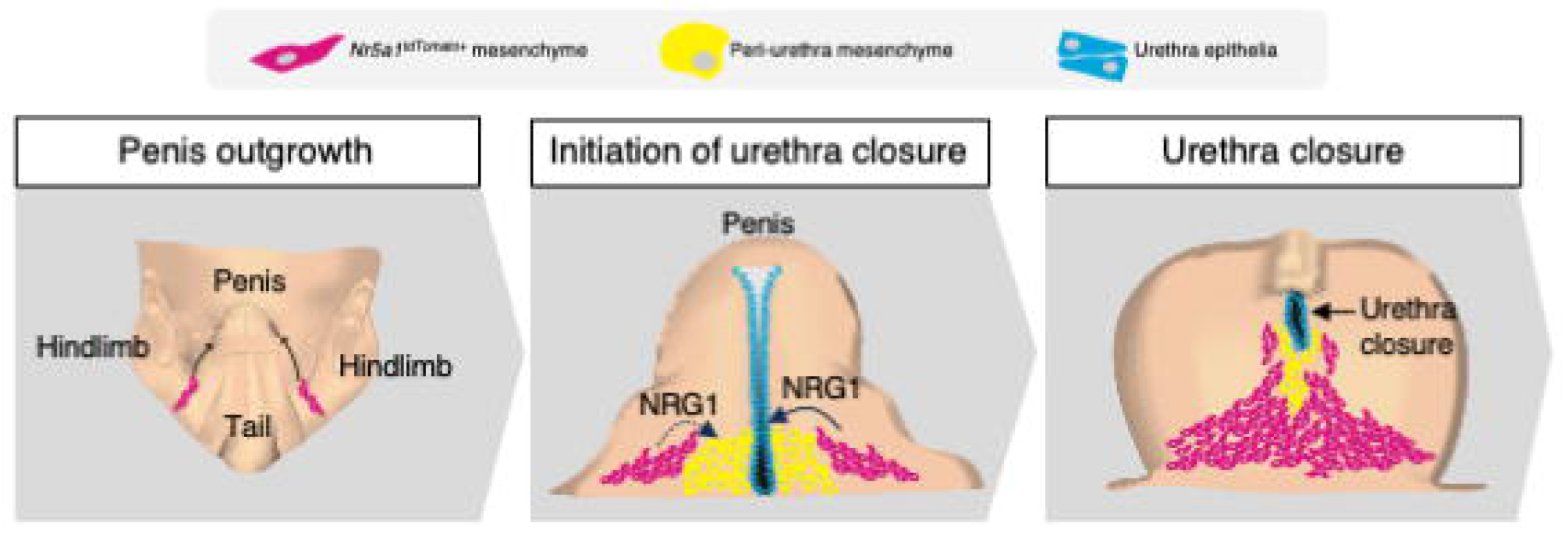
Summary of *Nr5a1^tdTomato+^* cell role in penis development. Just after penis outgrowth *Nr5a1^tdTomato+^* (magenta) cells are found posterior to the hindlimbs of the developing penis. As development progresses, these cells are found within the penis and are found on either side of the peri-urethra (yellow) and urethra epithelial cells (blue). At this time, there is extensive ligand-receptor communication between the *Nr5a1^tdTomato^* cells and the peri-urethra and urethral epithelium. As urethra closure progresses, the *Nr5a1^tdTomato^* cells move medially and converge at the ventral midline of the penis.

The dorsal portion of the penis is largely composed of cells derived from the umbilical region of the embryo (31). During tubercle formation, the umbilical derived cells migrate from the umbilicus into the dorsal portion of the penis (31). Surgical removal of umbilical derived cells caused hypoplasia of the dorsal penis and bladder exstrophy in mouse embryos (31). Despite the absence of umbilical derived cells, the ventral aspect of the penises still developed normally, and urethra closure was properly executed. The ventral portion of the penis is mostly composed of cells derived from the tailgut mesenchyme. Genetic ablation of critical genes in the tailgut mesenchyme, *Six1* and *Eya1*, caused severe hypoplasia of the ventral penis and severe urethra closure defects (32). In both the umbilicus and tailgut mouse models, the lateral aspects of the penis remained intact.

Cells derived from the inguinal region have been suggested to be involved in penis development through evolutionary studies and correlations of human genital defects with hindlimb abnormalities. In chickens, cells posterior to the hindlimb had the potential to occupy the genital tubercle (33). The hemipenes in squamate reptiles, which is a paired genital organ, form immediately adjacent to the hindlimb buds of the embryo (34, 35). Hypospadias and other genital defects in humans were correlated with hindlimb abnormalities. For example, cases of sirenomelia, a birth defect in humans where the hindlimbs fuse together during development, are significantly associated with hypospadias and penile hypoplasia (36, 37). In mice, loss of *Bmp4, Wnt5a,* or *Hoxa13* caused disruptions in both hindlimb and penis development (36, 38, 39). This circumstantial evidence pointed to the potential contribution of extra-genital cells near the hindlimb to penis formation. However, this hypothesis has never been definitively tested in a mammalian organism. We discovered that in the mouse embryo, the extra- genital cells originate near the inguinal region of the embryo and then migrate to the developing penis.

### The extra-genital cells provide essential paracrine factors for urethra closure

Cell-cell signaling is critical for normal urethra closure. Signaling pathways induced by FGFs, WNTs, BMPs, and SHH play distinct roles in shaping the fetal penis (10, 24, 25, 40, 41). Ablation of any of these signaling cascades caused severe penile defects. We found that the extra-genital cells, but not other cell population in the penis, have enriched express a ligand of the EGF signaling family, *Nrg1*. NRG1 is commonly associated with cell migration in Schwann cells and neural crest cells (30). In intestinal organoids of epithelial cells, NRG1 treatment induced actin-cytoskeleton alternations, which caused increased budding and tube formation (27, 42). NRG1 signals mainly through receptors ERBB3 and ERBB4, which commonly heterodimerize with EGFR *and* ERBB2 to facilitate signaling (43). *Egfr* and *Erbb3* were expressed in the peri-urethra cells and adjacent urethra epithelium, suggesting that NRG1 from the the extra-genital cells could act as a paracrine factor on either the peri-urethra cells or urethra epithelium to aid in urethra closure.

We found that when the extra-genital cells were ablated in the XY mouse embryos, hypoplasia of the prepuce and severe urethra closure defects occurred. These phenotypes resembled urethra epithelium knockout of β-catenin (44), which developed urethra closure defects but maintained the overall structural integrity of the genitalia. β-catenin is the effector molecule in response to WNT signaling (45). WNT and EGF signaling are tightly intertwined with one another in cancer, gut development, and cultured mesenchymal stem cells (46). Ligand bound EGF receptors can phosphorylate β-catenin in a similar way that WNT receptors induce β-catenin dependent gene expression (47). We suspect that these two signaling pathways coordinate to facilitate β-catenin dependent gene expression in the urethra epithelium, which is required for urethra closure.

### *Nr5a1* serves as a lineage marker for the extra-genital cells

NR5A1 is an important regulator of steroidogenic enzymes in the adrenal, pituitary, and gonads (48, 49). Discovery of *Nr5a1* expressing cells in the penis has never been documented. This finding led us to hypothesize that *Nr5a1* provides the developing penis with the unique capacity to metabolize testosterone into dihydrotestosterone. There are two main routes for dihydrotestosterone synthesis, one is through the canonical 5-α reductase (SRD5A2) pathway and the other is the backdoor dihydrotestosterone synthesis pathway which requires *Akr1c2* (50, 51). Both SRD5A2 and AKR1C2 are express in the embryonic human penis (51). *Nr5a1* moderately regulates *Srd5a2* and *Akr1c2* in prostate cancer cell lines (52). In the single cell sequencing data, *Akr1c2* and several other steroidogenic enzymes were not detected in the fetal mouse penis, therefore the backdoor pathway is unlikely. On the other hand, *Srd5a2* is detected in both the *Nr5a1^tdtomato+^* and *Nr5a1^tdtomato-^* cell populations of the penis. We found that in the *Nr5a1* knockout model, the penis developed normally and *Srd5a2* expression was not altered, indicating the NR5A1 is not responsible for inducing *Srd5a2* in the mouse penis. Based on the observation that *Srd5a2* expression is enriched in the male external genitalia compared to the female counterpart, it was speculated that either androgen signaling, or other male-specific genes could induce *Srd5a2* in the penis (13). In the liver, sterol regulatory element binding proteins (SREBF1/2) are clear modulators of *Srd5a2* expression (53). In our previous single cell sequencing data, *Srbef1* and *Srbef2* are ubiquitously expressed in the penis. They could be a potential regulator for *Srd5a2* during penile development.

Although *Nr5a1* expression in the penis has no functional role in urethra closure, *Nr5a1* marks a specific group of cells just after the onset of penis development. We found the expression of *Nr5a1* to be transitory, only lasting for one day, from E12.5- E13.5. The transitory nature of *Nr5a1* expression makes for a useful genetic tool to investigate the contribution of a small subpopulation of extra-genital cells in penis development. Tracing these cells through development have led to valuable insights into the origins of the penis, with future studies further teasing apart the direct contributions of these cell to penis development.

Despite that birth defects of the male urogenital system are some of the most common birth defects in the world, a large proportion of human hypospadias cases remain unexplained. This is likely due to the lack of information on gene regulation and cell population function within the developing penis. The research detailed above further expands our knowledge of basic penis development by identifying the functional importance of a unique cell population derived from extra-genital source, and their involvement in urethra closure by secreted EGF related ligands. With these insights the genetic and environmental contributors to hypospadias can be better identified, thus allowing us to develop preventatives or better alternative treatment for this far too common birth defect.

## Materials and Methods

### Mouse models

*Nr5a1-Cre^Tg/Tg^* (B6D2-*g(Nr5a1-cre)2Klp*) mice were generated by the late Dr. Keith Parker (54). *Rosa-LSL-tdTomato^f/f^* (B6.Cg-*Gt(ROSA)26Sor ^tm9(CAG-tdTomato)Hze^)*, Isl1- Cre^cre/+^ (Isl1tm1(cre)Sev/J), and *Rosa-LSL-DTA^f/f^* (C.129P2(B6)- Gt(ROSA)26Sortm1(DTA)Lky/J) were purchased at Jackson Laboratories (55). All mice used for lineage tracing and single cell RNA sequencing contain the alleles for *Nr5a1- Cre^Tg/+^;Rosa-LSL-tdTomato^+/f^. Nr5a1-Cre^Tg/+^; Rosa-LSL-tdTomato^+/f^* were generated by crossing *Nr5a1-Cre^Tg/Tg^* male mice with female *Rosa-LSL-tdTomato^f/f^*. Observation of a vaginal plug was defined as embryonic day or E0.5. At dissection developmental stage was confirmed by the Theiler staging criteria, and mice were sexed by evaluating presence of testis or ovaries. Presence of tdTomato fluorescence was used to confirm the genotype.

*Nr5a1^tdTomato+^* cell ablation experiments used a mouse model with a diphtheria toxin gene inserted into the Rosa allele (*Rosa-LSL-DTA^f/f^*) (21). To generate *Nr5a1^tdTomato+^* cell ablation mice (*Nr5a1-Cre^Tg/+^; Rosa-LSL-tdTomato^+/f^; Rosa-LSL-DTA^+/f^*), *Nr5a1-Cre^Tg/+^; Rosa-LSL-tdTomato^+/f^* male mice were crossed with *Rosa-LSL- DTA^f/f^* female mice. Embryos were sexed by the presence (female) or absence (male) of the Barr body via Giemsa staining of the amnion (56) and were genotyped by probing for the presence of a Cre allele. We confirmed that *Nr5a1-Cre* also targets the adrenal glands and activation of diphtheria toxin caused a complete loss of adrenals **(Supplemental 6).** To determine the role of lack of androgens in the phenotype of these *Nr5a1^tdTomato+^* cell ablation embryos, pregnant *Rosa-LSL-DTA^f/f^*female were supplemented daily subcutaneously with 0.2 mg/10g testosterone propionate (Sigma Aldrich T1875-5G) from E13.5 to E18.5 to replace lost testosterone due to testis ablation. All animal procedures were approved by the National Institute of Environmental Health Sciences (NIEHS) Animal Care and Use Committee and follow a NIEHS-approved animal study proposal.

### Slice culture of the external genitalia

*Rosa-LSL-tdTomato^f/f^* female mice crossed with *Nr5a1-Cre^Tg/Tg^*male mice were euthanized at E15.5. Once pups were removed from the uterus and decapitated, bodies were kept on ice. Male embryos were separated based on the presence of testis. Female embryos were not used in these studies. Once sex was defined, male genitalia were removed and embedded in 4% low melting agarose (GeneMate E-3112-125). Low melting agarose was mixed with sterile 1X PBS and microwaved for 1 minute to dissolve, and it was kept at 37C for a minimum of one hour before embedding. Leica VT1000S vibratome was used to make 150 µm coronal sections of penises. Section speed was between 0.1mm/s-0.3mm/s to maintain tissue integrity. All sectioning was conducted in ice cold, sterile 1X PBS. Sections were selected for culture based on 1) presence of tdTomato fluorescent *Nr5a1^tdTomato+^* cells and 2) lack of urethra closure. The selected sections were placed in a 0.4µm cell culture insert (Millipore PICM01250) that was saturated with slice culture media. Slice culture media consisted of phenol-red free DMEM /F12 (Gibco 21041-025), 10% charcoal stripped fetal bovine serum (Sigma Aldrich F6765), and 1% penicillin/streptomycin (Sigma Aldrich P4333).

Dihydrotestosterone (10^-4^ M) (Steraloids A2570-011) or Laptinib (1µM) (Sigma-Aldrich D8418-100ML) was added to examine the loss of EGF related signaling. Culture inserts were placed inside either a 35 mm glass bottom petri dish (Ted Pella 14022-1120) or a 12 well-glass bottom culture plate (Cellvis P12-1.5H-N) with 1 mL of slice culture media. Petri dish was placed into Keyence BZ-X810 for live cell imaging. Images were taken at 10-minute intervals with a 10 µm z-stack with a 10x objective. Imaging lasted for at least 48 hours to capture urethra closure. The captured images were compiled into movies with the Keyence BX-X800 analyzer. To create cell tracks, movies were imported into Imaris, and the Dragon tail feature was applied to red fluorescing cells.

### RNAscope, immunofluorescence, and light-sheet imaging

Penises were fixed in 4% paraformaldehyde overnight at 4C, watched in 20% sucrose in 1XPBS, embedded in optimal cutting temperature compound and sectioned at 10µm. Sections were mounted on Superfrost Plus slides (Fisherbrand, 12–550-15) and kept at -80C until further processing. For RNAscope probes *Scube2* (ACD 488141)*, Nrg1* (ACD 418181-C3)*, Erbb3* (ACD 441801-C2)*, Grem1* (ACD 314741-C3)*, tdTomato* (ACD 317041 and 317041-C2), and *Srd5a2* (ACD 431361). Were used according the ACDBio’s protocol. For immunofluorescence, sections were permeabilized in 1xPBS with 0.1% Triton and blocked for 1 h with 10% donkey serum. Primary antibodies against AKAP12 (gift from Irwin H. Gelman, Roswell Park Cancer Institute), dsRED (Clontech 632496), FOXL2 (Abcam ab5096), EGFR (Abcam ab32077) and FOXA1 (Abcam ab170933) (Supplemental table 2) were incubated on slides at 4C overnight. After incubations, slides were washed with 1%Triton, and secondary (Alexa Fluor 488, 594, and 647) were incubated for 2 h at room temperature. Slides were counter-stained with DAPI (1:1000)(Millipore 5.08741.0001), mounted with Pro-Long Antifade (Invitrogen P36970), and were imaged on a Zeiss 900 confocal microscope the day after processing.

For lightsheet imaging, samples were fixed in in 4% paraformaldehyde overnight at 4C and then cleared by strictly using the iDISCO protocol (57). Briefly samples were dehydrate with a methanol series, then bleached with 3% H_2_O_2,_ delipidated with 66% dichloromethane, and rehydrated with a methanol series. Samples were then permeabilized, blocked, and labeled with dsRED (Clontech 632496) and a anti-rabbit alexafluor 647 secondary antibody. Finally, the samples were dehydrated in a methanol series, delipidated with dicholormethane, and cleared in dibenzyl ether. Samples were image on the LaVision ultramicroscope II at University of North Carolina-Chapel Hill. Images were processed with Imaris.

### Cell disassociation and single-cell suspension

Preparation for single cell sequencing follows the same protocol in our published paper (18). Penises from E16.5 embryos (n=2) were placed in 250µL of ice-cold disassociation media (sterilized 1x PBS and 0.04% bovine serum albumin, 1.2 U/mL Dispase II (Roche 04942078001), 1 mg/mL Collagenase IV (Gibco 17104–019), and 5 U/mL deoxyribonuclease I (Roche 1010459001)). After the tissues were successfully homogenized, the cells were washed with 1x PBS with 0.04% bovine serum albumin. Cells were loaded into the 10x Chromium machine for single cell separation.

### Library preparation, sequencing, and sequence alignments

Data from GSE174712 was used for the initial characterization of the *Nr5a1^tdTomato+^* cells (18). For the cell ablation model, library preparation was conducted following the 10x Genomics library preparation protocol. The libraries were pooled into a single tube and sequenced on an Illumina NovaSeq 6000 with an S2 flow cell. Paired-end sequencing was conducted with the first read being 26-bp (base pair) and the second read being 96-bp. DTA libraries were sequenced to at least 13,000 total reads per cell (**Supplemental Table 1**), which was sufficient to obtain clustering and differential gene expression analysis. Both library preparation and sequencing were performed by the Epigenomics and DNA Sequencing Core Facility at NIEHS. Align sequences. To detect *tdTomato* mRNA transcripts in the single cell sequencing data, we added the *tdTomato* coding sequence (Supplemental data file 1) to the mm10 genome and genes files. Cellranger v3 was then used to align the data with the *tdTomato* sequence.

### Single cell data analysis

Aligned and annotated data sets were imported into R for Seurat analysis (v. 4.2.1) (58). Cells with < 7500 and > 1000 mRNA counts were removed from the data set. Cells with mitochondrial content > 25%, Hba content > 25%, and Hbb content >50% were removed from the data sets. Log normalization, data scaling, dimensionality reduction and clustering was conducted both the cell ablation and developmental series data sets (Rcode 1 and 2). Once data were processed, we tested whether *tdTomato* could be properly detected. After plotting a UMAP of *tdTomato* gene expression, it was apparent that the transcript expression was leaky as previously reported by (59), although leaky transcript expression did not result in red fluorescent protein in all genitalia cells. There was a low background level of expression throughout most cells.

Although most cells did express *tdTomato*, there was a clear separation of high or low tdTomato expressing cells. Thus, cells with <10 counts of tdTomato expression were considered *Nr5a1^tdTomato^*negative and cells with >10 counts were *Nr5a1^tdTomato^* positive. To validate that *Nr5a1^tdTomato^* positive cells truly represented the cells identified within the mouse penis, we identified genes that were significantly enriched in the *Nr5a1^tdTomato^*positive cells (Dataset 1) and conducted RNAscope for markers *Grem1*, *Scube2,* and *Nrg1.* Once *Nr5a1^tdTomato^* positive cells were correctly identified, the developmental series dataset was used to investigate ligand-receptor interactions. Epithelial, peri- urethra, and *Nr5a1^tdtomato+^*cells were subsetted into a separate dataset and ligand receptor analysis was executed with Cellphone DB v2.0 ((26) and Data set 2).

The cell ablation dataset was processed the same as the developmental series data. To identify differential expression between control and *Nr5a1^tdTomato^* cell ablated mice, likelihood ratio tests were conducted between the genotypes for each cell population (Rcode). To further investigate the changes within the peri-urethra, that cell population was subsetted reclustered, and visualized with a UMAP.

### Statistical analysis

For individual gene expression analysis, anogenital measurement comparisons, and hypospadias severity comparisons linear mixed effect models were used to calculate statistical differences (p<0.05) (60). For single cell gene expression analysis, each biological replicate was treated as the random effect. For anogenital distance and hypospadias severity comparisons, dam was classified was the random effect.

## Acknowledgements

We thank Dr. Meilan Zhao for training on the vibratome for slice cultures. We also thank Dr. Pablo Ariel for help with light sheet imaging and Dr. Blanche Capel and Corey Bunce for providing *Nr5a1-Gfp* mice. We are grateful to the NIEHS Epigenomics Core for the single cell sequencing, Integrative Bioinformatics Support group for support when analyzing single cell data, and Comparative Medicine Branch for mouse colony maintenance. This work was supported by the Intramural Research Program (ES102965 to H.H.-C.Y.) of the NIH, National Institute of Environmental Health Sciences and the K99/R00 fellowship (1K99DK132460-01A1 to C.M.A.) from the National Institute of Diabetes and Digestive and Kidney Diseases .

---

**Supplemental Figure 1.**
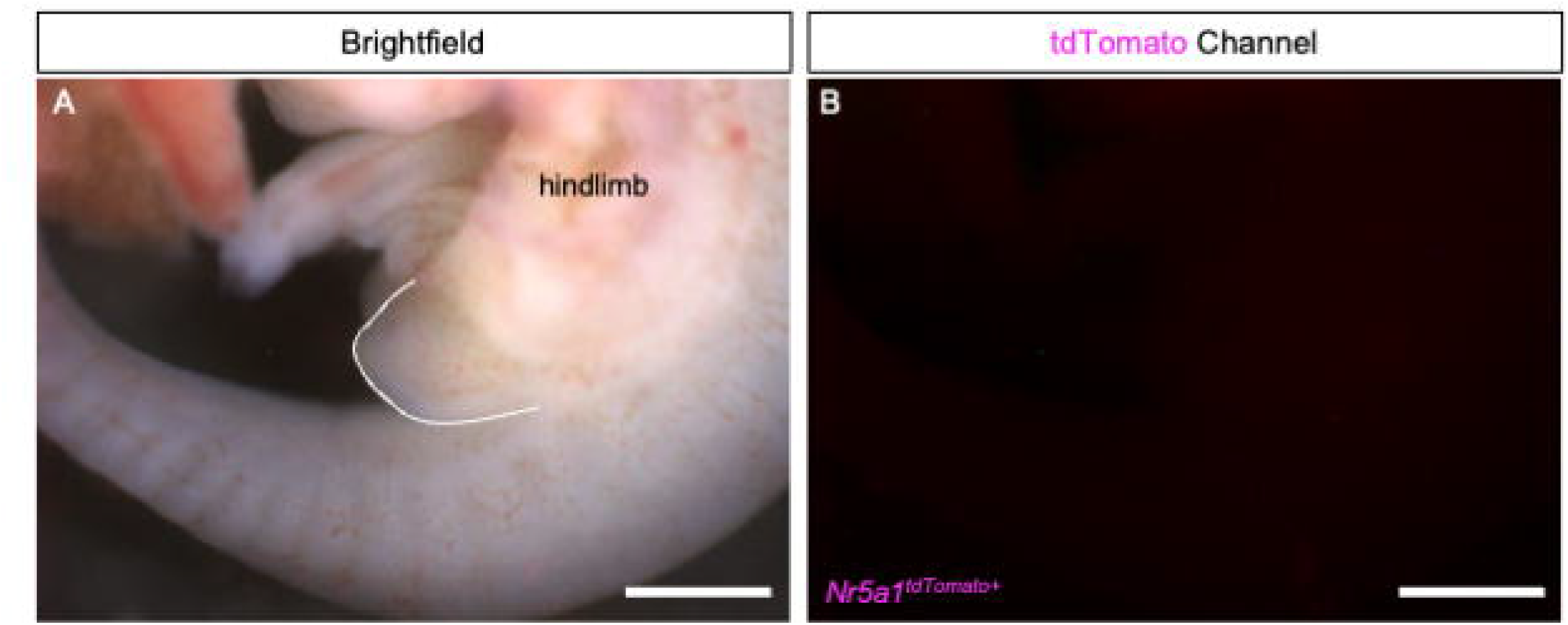

**Supplemental Figure 2.**
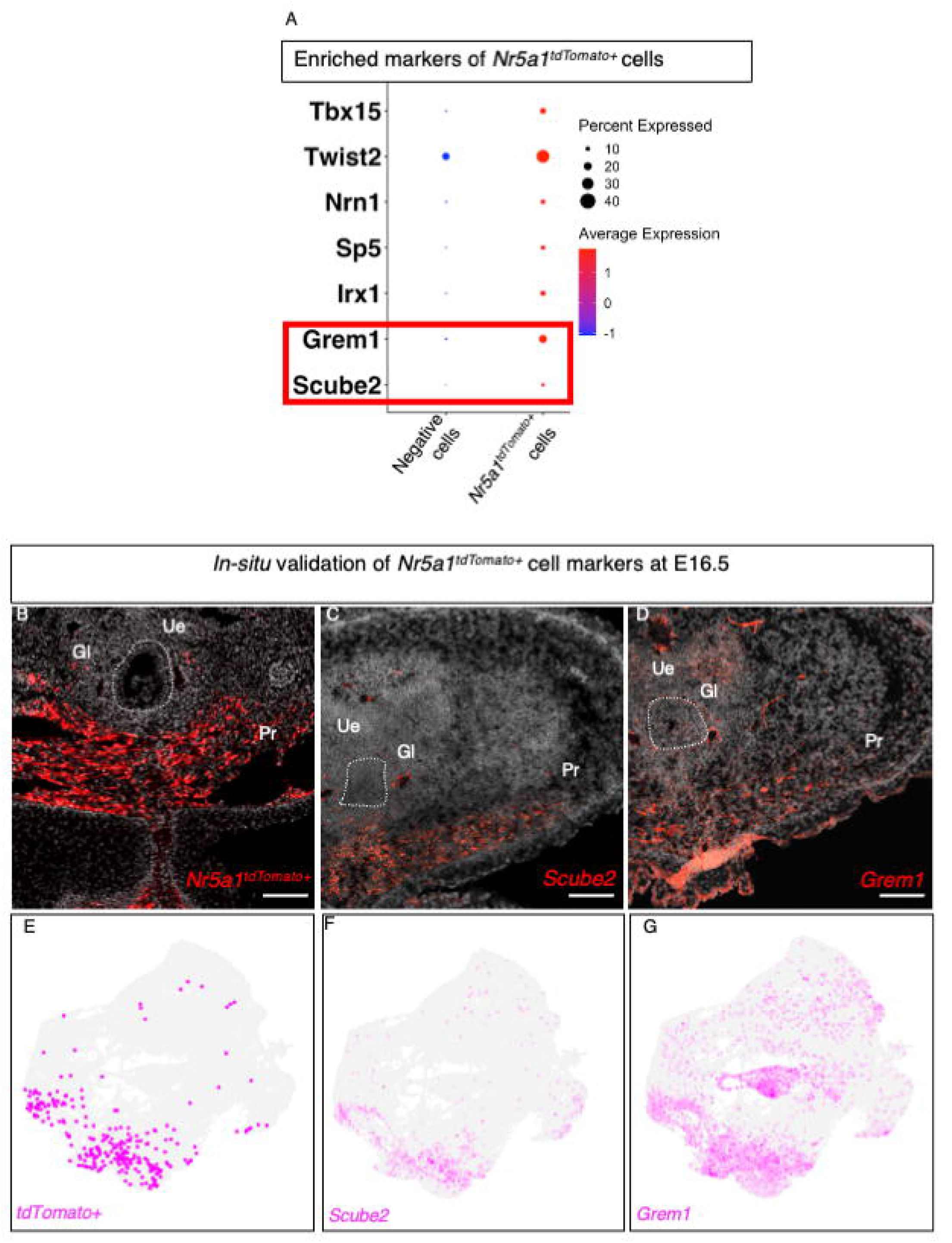

**Supplemental Figure 3.**
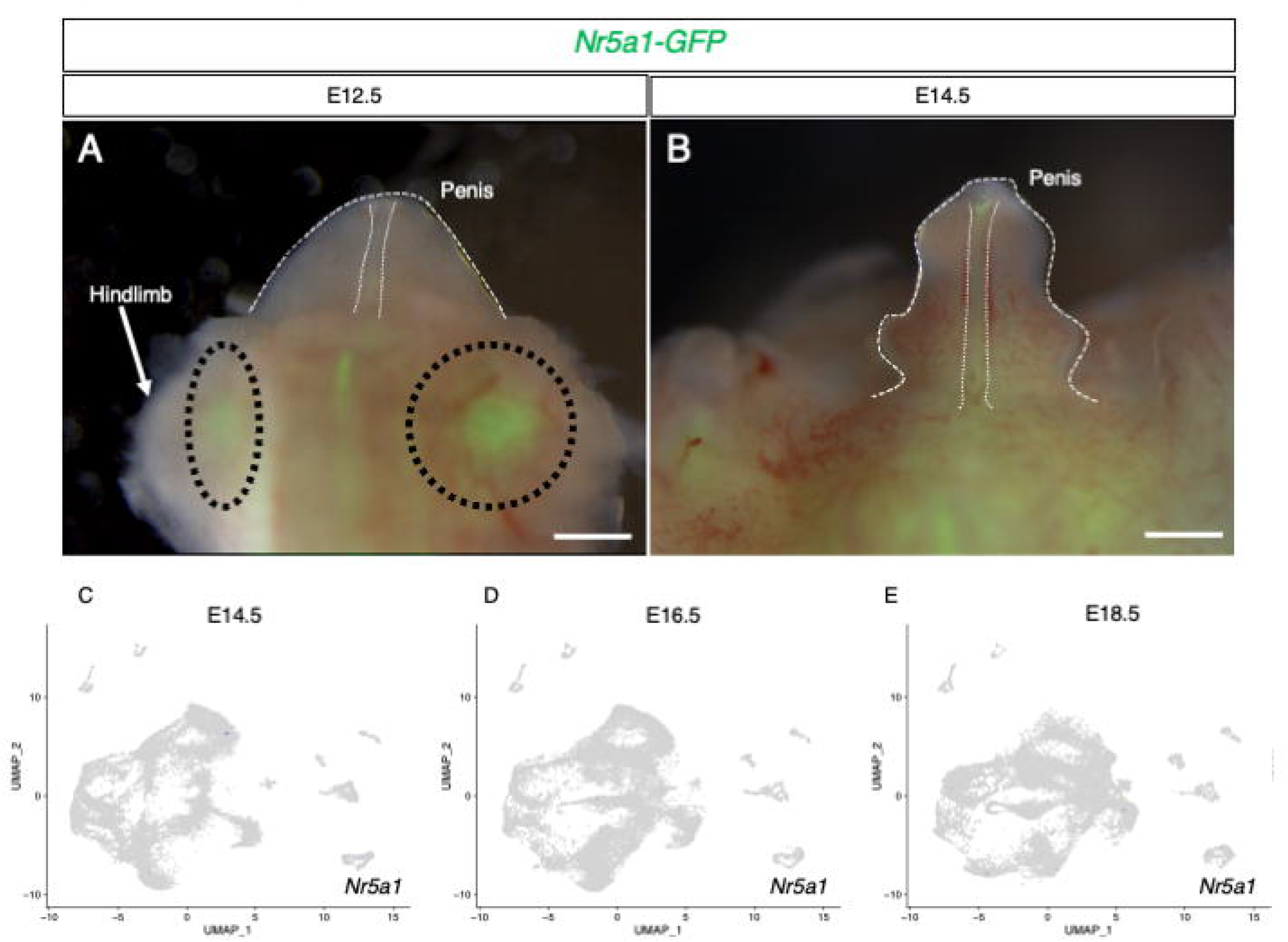

**Supplemental Figure 5.**
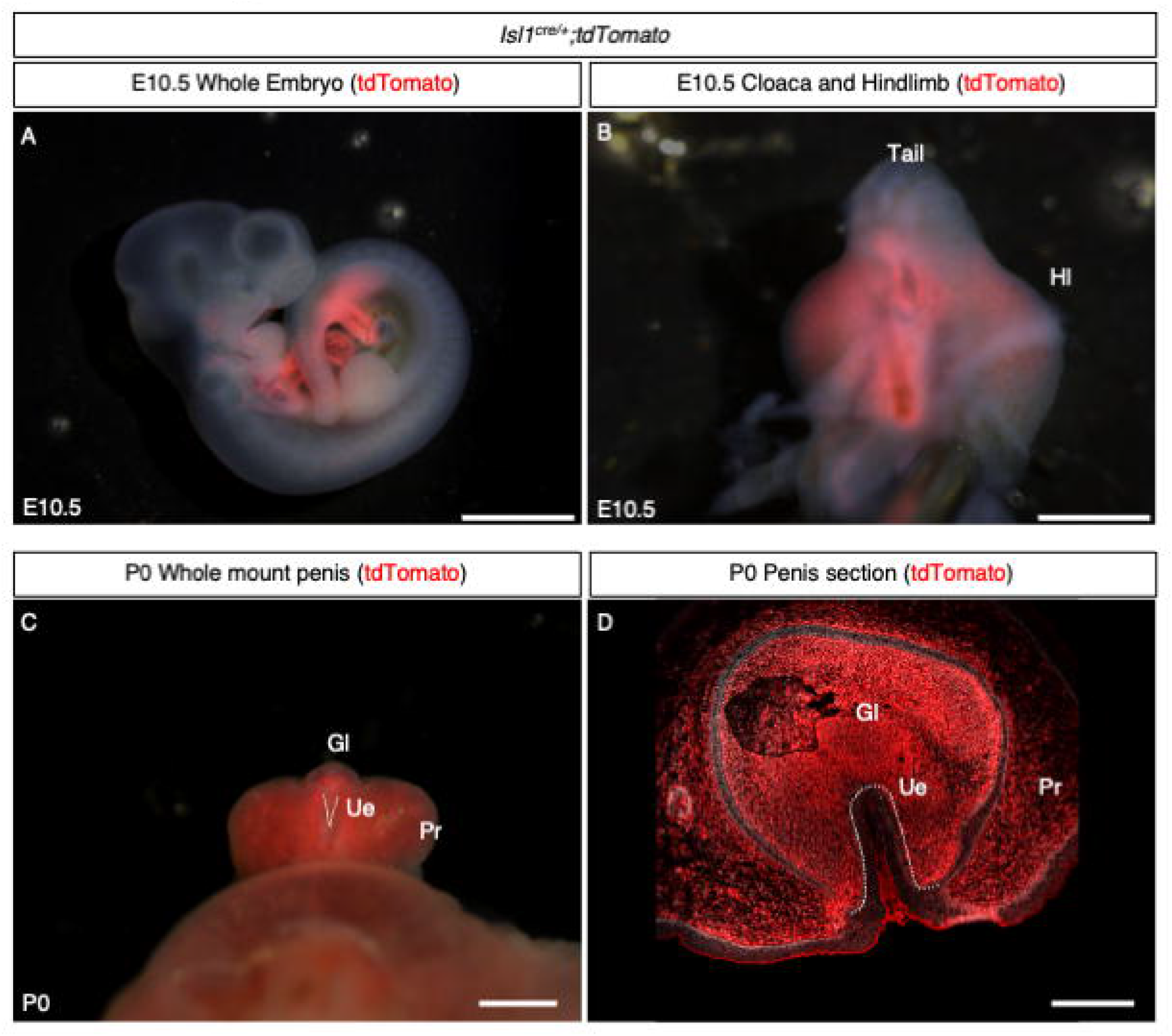

**Supplemental Figure 5.**
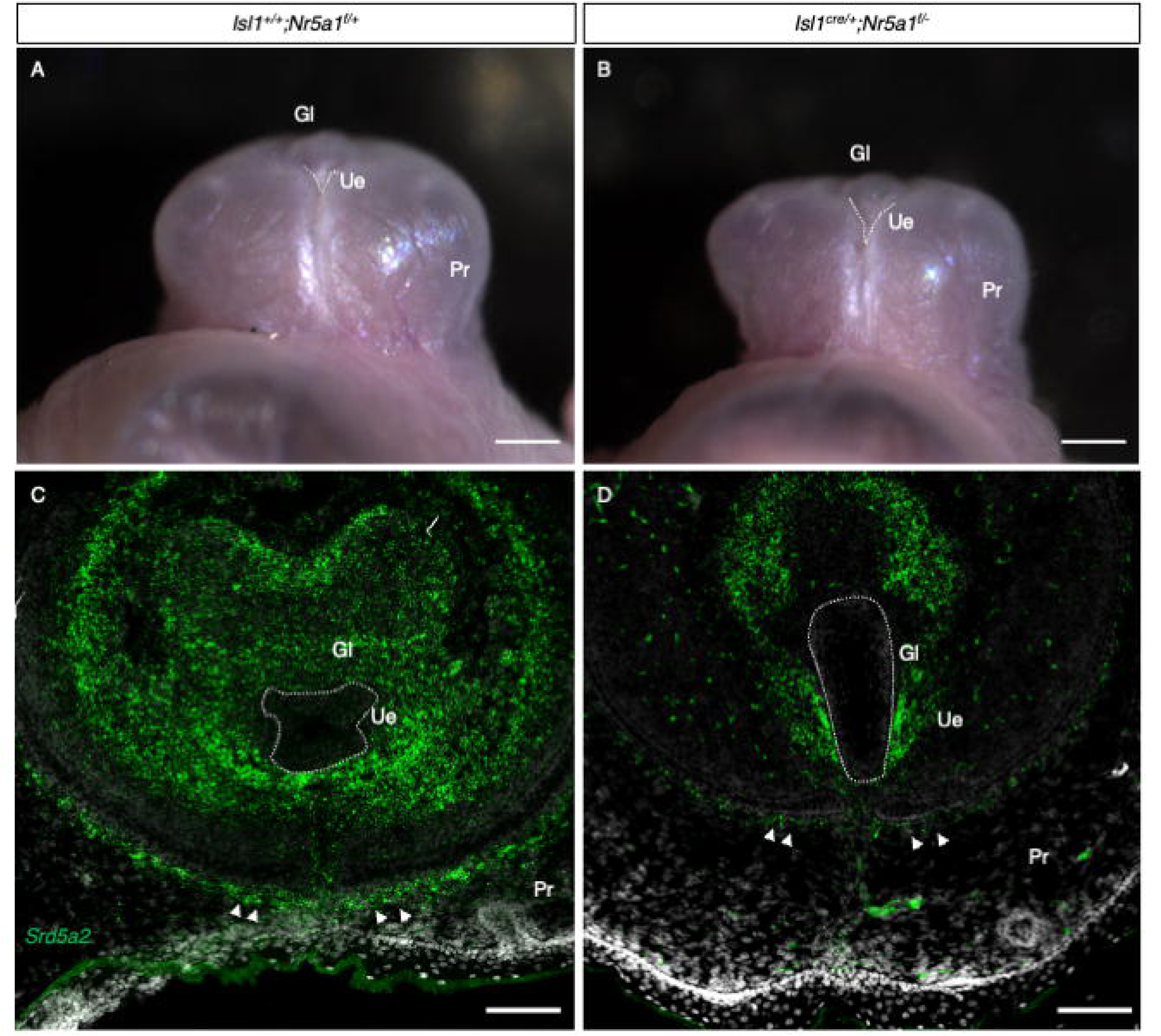

**Supplemental Figure 6.**
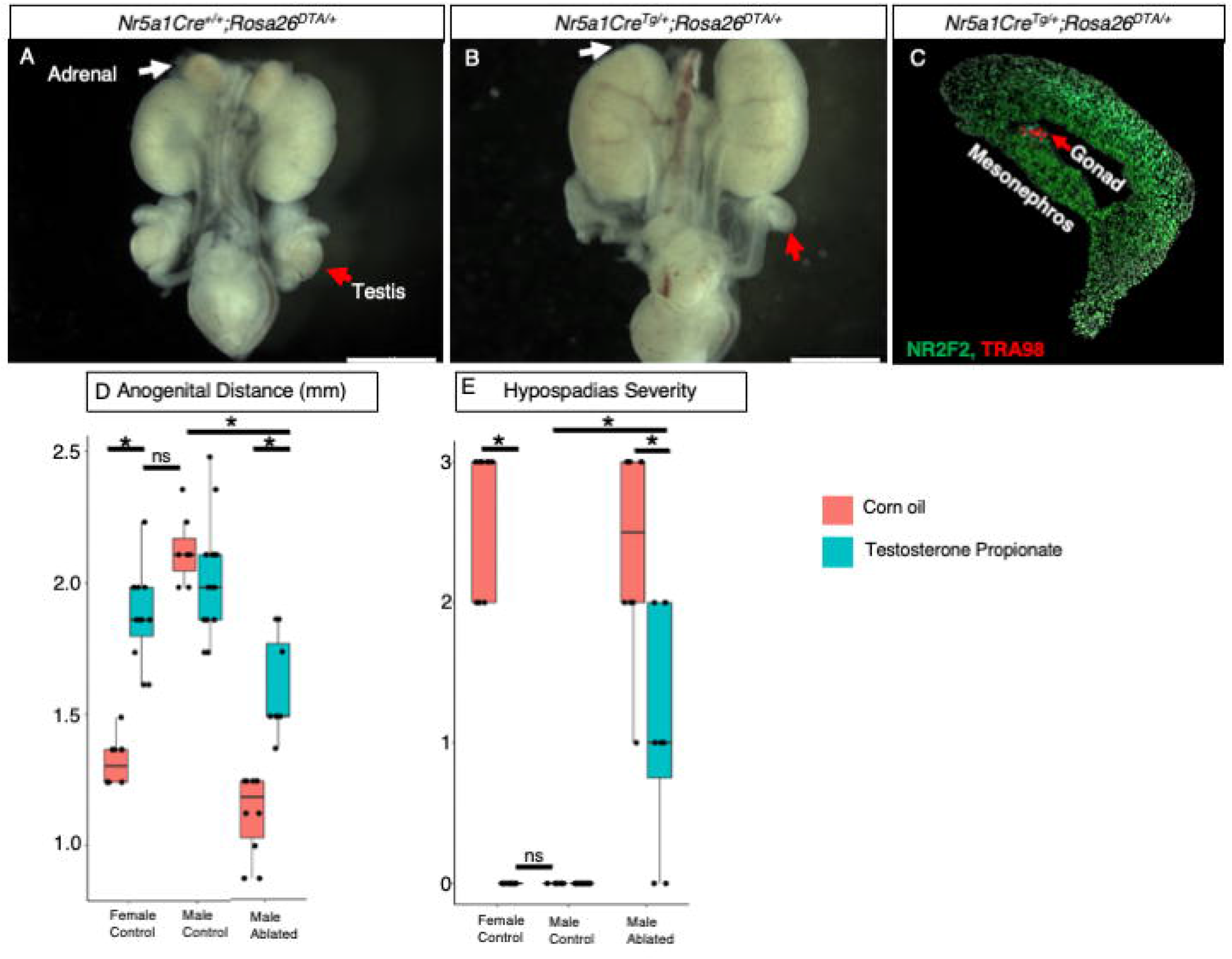

**Supplemental Figure 7.**
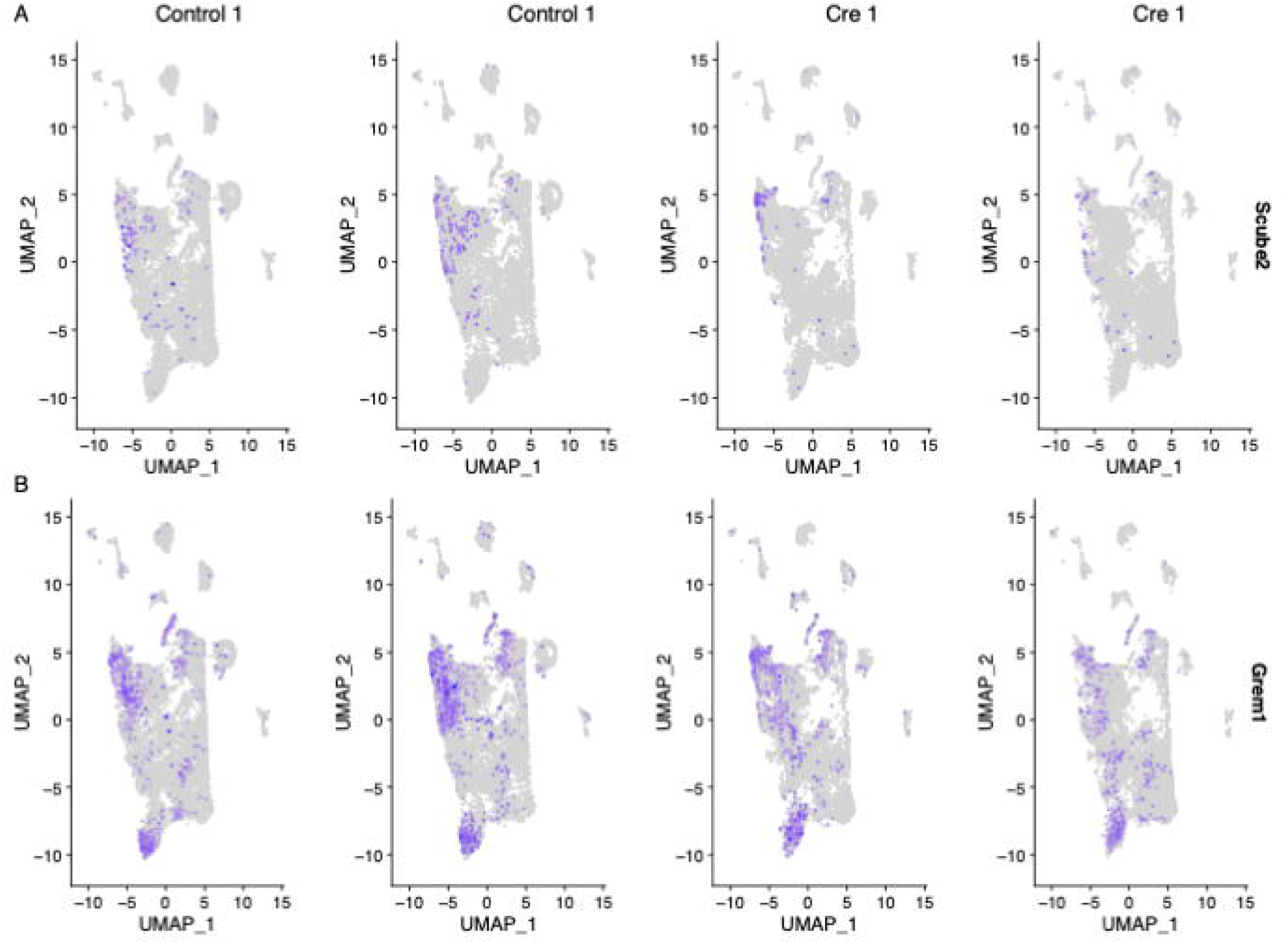

**Supplemental Figure 8.**
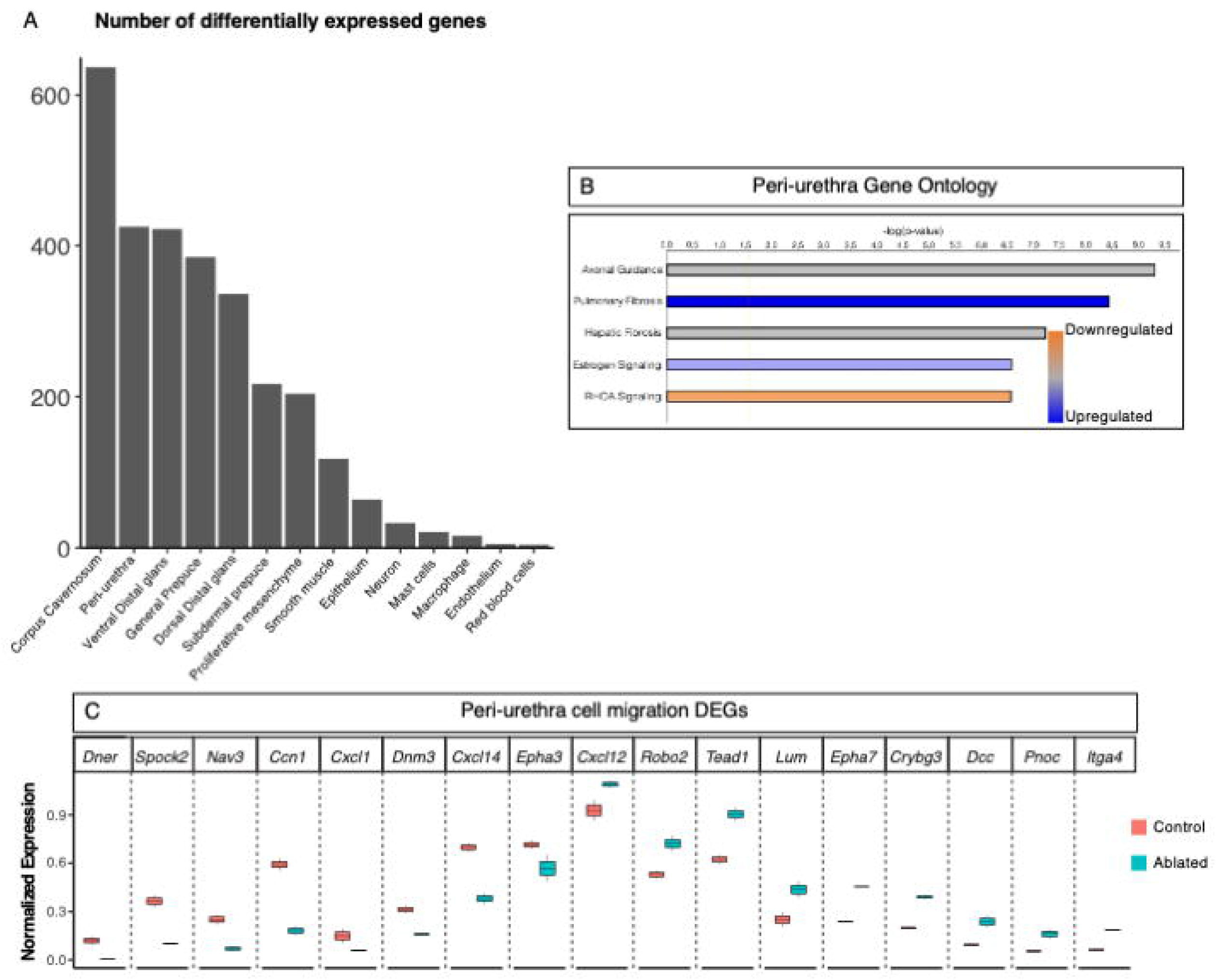

