## Supplemental table 1 for "An extra-genital cell population contributes to urethra closure during mouse penis development"

Supplemental Table 1. Sequencing results from *Nr5a1*$^{tdtomato+}$ cell ablated single cell

| Sample | Raw.Reads | Barcode | Mapped | Saturation | Cells | Reads/ Cell | Median umi/cell | Median genes/cell |
|---|---|---|---|---|---|---|---|---|
| DTA 1 | 163,797,148 | 93% | 87% | 27% | 11,511 | 14,230 | 3,578 | 1,478 |
| DTA 2 | 159,515,741 | 96% | 89% | 28% | 11,349 | 14,055 | 3,661 | 1,592 |
| MALE 1 | 221,873,673 | 97% | 86% | 35% | 12,570 | 17,651 | 4,940 | 1,816 |
| MALE 2 | 153,731,758 | 96% | 87% | 25% | 11,613 | 13,238 | 3,992 | 1,627 |
