## Supplemental Table 2 for "An extra-genital cell population contributes to urethra closure during mouse penis development"

Supplemental Table 2. Antibodies

| Target | Company | Catalog number | Concentration |
|---|---|---|---|
| AKAP12 | Collaborator | N/A | 1:300 |
| FOXL2 | Abcam | Ab5096 | 1:300 |
| EGFR | EGFR | ab32077 | 1:400 |
| FOXA1 | FOXA1 | Ab170933 | 1:200 |
| dsRED | Clontech | 632496 | 1:1000 |
| E-cadherin | Abcam | Ab11512 | 1:500 |
| Alexafluor 488 | Invitrogen | A21208/A21206 | 1:400 |
| Alexfluor 568 | Invitrogen | A10042 | 1:400 |
| Alexfluor 647 | Life Technology | A21084 | 1:400 |
