## Supplemental Figure Legends for "An extra-genital cell population contributes to urethra closure during mouse penis development"

**Supplemental Movies**

Supplemental movie 1. **Lightsheet z-stack of *Nr5a1$^{tdTomato+}$* cells in E12.5 embryo**

Supplemental movie 2. ***Nr5a1$^{tdTomato+}$* cell migration during urethra closure**

Supplemental movie 3. ***Nr5a1$^{tdTomato+}$* cell migration during urethra closure in control slice**

Supplemental movie 3. ***Nr5a1$^{tdTomato+}$* cell migration during urethra closure in lapatinib exposed slice**

**Supplemental Figure Legends**

**Supplemental Figure 1. *Nr5a1$^{tdTomato+}$* cells are not found near hindlimbs at E11.5**. (A) Brightfield images of E11.5 embryo with the solid white line outlining the genitalia. (B) Fluorescent image of the same E11.5 embryo. Scale bar = 500 μm

**Supplemental Figure 2. Other markers *Nr5a1$^{dTomato+}$* cell population.** (A) Dotplot of significant *Nr5a1$^{tdTomato+}$* cell markers with the size of the dot representing the percentage of cells expressing the gene and the color of dot showing relative expression (red=high and blue = low). (B-D) Immunofluorescence of *Nr5a1$^{tdTomato+}$* cells, *Scube2,* and *Grem1* in penis sections at E16.5. (E-G) UMAP of *tdTomato*, *Scube* gene expression in E16.5 penis. (D) *Scube2* RNAscope in E16.5 penis and (E) UMAP gene expression of *Scube2* from single cell mRNA sequencing data. (F) Grem1 RNA scope in E16.5 penis and (G) UMAP gene expression of *Grem1.* P= penis, Gl = Glans, Ue = urethra, and Pr= Prepuce. Scale bars= 50μm

**Supplemental Figure 3. Investigating active expression of *Nr5a1* in penis development**. (A and B) Lower half of *Nr5a1-GFP* mouse embryo at E12.5 and E14.5. White solid line outlines the genitalia. Black dotted circles outline the active expressing regions of GFP in the E12.5 embryo. (C-E) Single cell mRNA sequencing gene expression UMAPs for *Nr5a1* with grey dots displaying no expression and dark blue showing high expression. Scale bars = 800µm

**Supplemental figure 4. Validation that *Isl1Cre* targets the external genitalia prior to genitalia formation.** (**A** and **B**) Endogenous fluorescence of tdTomato at E10.5. **(C)** Whole mount image of external genitalia at P0 with endogenous red fluorescence. (**D**) histological section of P0 genitalia with red representing endogenous fluorescence. Hl=Hindlimb, Gl = Glans, Ue = urethra, and Pr= Prepuce. Scale bar for A = 1.4 mm, B = 540 µm, C = 800 µm, and D = 100µm

**Supplemental Figure 5. Knockout of *Nr5a1*, the gene, in the penis. (A and B)** Whole mount images of control (A) and knockout (B) mouse penises. White dotted line represents the open urethra. (**C** and **D**) RNAscope for *Srd5a2* on penis sections with the dotted line representing the urethra epithelium and the white arrow indicting predicted regions of $Nr5a1^{tdtomato+}$ cell localization. Gl = Glans, Ue = urethra, and Pr= Prepuce. Whole mount scale bars = 200 µm and microscope scale bars = 100 µm

**Supplemental Figure 6. Investigation of *Nr5a1* cell ablation in other organs. (A and B)** Whole mount image of the urogenital complex of control (“exact name in figure”)

and ***Nr5a1* cell** ablated mice. White arrow indicates positioning of the adrenal and red arrow indicated positioning of the testis. (**C)** Immunofluorescent image of a *Nr5a1*$^{tdtomato+}$ cell ablated gonad with red labeling TRA98+ germ cells and green labeling NR2F2+ interstitial cells of the mesonephros (and a few from the gonad I suppose). The red arrow designates the gonad. (**D**) Anogenital distance length for control and *Nr5a1*$^{tdtomato+}$ cell ablated mice exposed to either corn oil (red) or testosterone propionate (teal). Asterisk indicate p<0.05. (**E**) Hypospadias severity scoring for control and *Nr5a1*$^{tdtomato+}$ cell ablated mice exposed to either corn oil (red) or testosterone propionate (teal). A score of 3 represents severe hypospadias and a score of 0 represents no hypospadias. Asterisk indicate p<0.05. Whole mount scale bar = 2mm

**Supplemental Figure 7. Expression of *Nr5a1*$^{tdTomato+}$ enriched genes.** (**A**) Expression of *Scube2* in control and Cre+ (*Nr5a1*$^{tdTomato+}$ cell ablated) mice. (**B**) Expression of *Grem1* in control and Cre+ mice.

**Supplemental Figure 8. Impacts of *Nr5a1* cell ablation on other penis cell populations.** (**A**) Differential gene expression between wildtype and *Nr5a1*$^{tdtomato+}$ cell ablated mice for each penis cell population. (**B and C**) Gene ontology of the differential expressed genes found in the peri-urethra and prepuce. Grey bars represent no discernable up or down regulation of the pathway, increasingly red bars indicate elevation of the pathway. (**D** and **E**) Gene expression boxplots of control (red) and ablated (teal) for significantly altered genes in the prepuce (D) and peri-urethra (E).
